# A pump-leak model of epithelia reveals the roles of paracellular pathways in water transport & the importance of boundary conditions

**DOI:** 10.64898/2026.10.26.760817

**Authors:** Alan R. Kay, Zahra Aminzare

**Affiliations:** Depts. of Biology, and University of Iowa, Iowa City, IA 52242; Mathematics. University of Iowa, Iowa City, IA 52242; Dept. of Pediatrics, Section of Developmental Biology, University of Colorado Anschutz Medical Campus, Aurora, CO 80045

**Keywords:** Epithelia, water, sodium, potassium, chloride, pumps

## Abstract

We develop a pump-leak model of epithelial transport, where the flux of water through the epithelium is modeled as a compartment, or sink, whose volume grows linearly with time. A method is described for calculating the voltage in the sink compartment, as well as analytical solutions to the system of equations. We show how growing boundary conditions can be used to model a scenario where water and ion fluxes are possible in the steady state. It is possible to drive a continuous flux of water through an epithelium that is powered by the Na^+^ pump establishing small osmotic gradients between compartments. We show that paracellular pathways play a crucial role in the operation of epithelial transport by ensuring that the flux through the epithelium is electroneutral. For the Na^+^ pump situated on the basolateral membrane, water fluxes can be directed in the same direction as the Na^+^ ion by having a high Na^+^ conductance on the apical membrane and high K^+^ conductance on the basolateral membrane. Flipping these conductances changes the direction of water fluxes. We introduce a graphic formalism that makes clear the precise mechanism of water transfer, showing the osmotic and electrical constraints. We demonstrate that it is possible when the extracellular osmolarity is low (∼10mOsm) for the intracellular osmolarity to exceed that of the former considerably and produce a hyperosmolar solution.

## Introduction

Epithelia play an important role in the physiology of all animals, orchestrating the fluxes of ions, metabolites and water, through the polarized arrangement of transporters in their apical and basolateral membrane (Berridge and Oschman, 1972). The transport of water, which represents about 70% of a cell’s mass, is essential for the survival and growth of any cell in all the kingdoms of life (Dick, 1966; Potts and Parry, 1964). Even after many years of study, precisely what drives water across epithelia remains one of the central enigmas in biology (Caceres et al., 2017). There have been several theories put forward, but no definitive mechanism has yet been proposed that can account for water transport.

In this paper, we will show that the mechanism that is likely to underpin water transport is the pump-leak model (PLM), which was formulated by Tosteson and Hoffman (1960) to account for the stabilization of cell volume by the Na^+^/K^+^ ATPase (NKA). Surprisingly, although water transport lies at the heart of the PLM, it has with some exceptions (Hoffmann and Ussing, 1992) not been widely considered in the context of epithelial water transport.

Epithelial physiology is rife with diagramatic representations of cells with the various transporters and channels on the apical and basolateral membrane. However, although such diagrams are useful they do not and cannot provide a full understanding of how epithelial cells function. In this paper we show how one can model the transport of ions and water in an epithelial system, with a system of differential and algebraic equations that encapsulates the physico-chemical forces acting on the ions and water.

### Outline of the PLM

Textbooks often suggest that the role of the NKA is to establish an asymmetry in Na^+^ and K^+^ across the membrane to generate a negative membrane potential (Alberts et al., 2015). While this is certainly true, it has become clear that it is not the pumps primary function. Only once one lays out the essentials of the PLM, does its function become apparent. To do this, one has to consider the physical forces that act on ions and water across the membrane. One sets up a system of differential equations that encompases the forces acting on ions and water as well as the cells mechanics (see below). These pump-leak equations (*Development of the ABp equations*) encapsulate how ions and water distribute as a result of the action of the pump. They show that the operation of the pump stabilizes the cell against the osmotic flux of water which threatens its stability. Moreover, it drives Cl^-^ out of the cell making room for metabolites that are needed at adequate concentrations to power the cell (Aminzare and Kay, 2024). By this we mean that since the intracellular osmolarity is constrained to about 300 mOsm, with cations occupying about 150 mOsm, there is only room for ∼150 mOsm of anions. The generation of a negative membrane potential is simply a consequence of the action of the NKA.

The physical forces operating within the cell lead to the development of two constraints; an osmotic one that ensures that the osmolarities of the intracellular and extracellular solutions balance, and an electric constraint that ensures that in a given macroscopic cellular compartment, that there is rough electroneutrality. It should to be emphasized that these constraints emerge from the physics of the system and are not imposed externally.

### Some definitions

Water transport across an epithelium can occur in one of two directions: 1) **Absorptive** transport, from the apical to the basolateral side. 2) **Secretory** transport, from the basolateral side to the apical side. We will refer to the side from which the transported liquid arises as the **source** and the side into which it moves the **sink**. If the fluid flowing into the sink (i.e., transportate) has exactly the same osmolarity as the source, then the transport process is **isotonic** (sometimes isosmotic) (Tripathi and Boulpaep, 1989). In some cases, the source and the sink have almost the same ionic composition (**iso-ionic**).

### Isosmotic Transport

Much of the debate around epithelial water transport has centered around the notion of isotonic transport. However, as has been pointed out it was not and still is not possible to distinguish solutions which have an osmolarity difference less than about 1-3% (Spring, 1999). As we will show below, it is possible to drive water fluxes with minute differences in osmolarity (∼0.001%). We suggest that this mechanism should be termed **near-isosmotic** (Altenberg and Reuss, 2013). This is an important distinction since, as we will show, it is the difference in osmolarity that drives water transport; no transport would be possible if the osmolarity of the solutions flanking the epithelium were the same.

Our approach is to use as simple a model as possible, on the grounds of parsimony, to understand how water might be transported across an epithelium. There has been a tendency in the simulations of epithelial transport to try and build an accurate model of one type of epithelium, adding the appropriate channels and transporters, to see if it replicates the phenomena. Our approach is a little different in that we are looking for overall principles of epithelial transport.

We adopt what we will call an osmo-electrodiffusion approach, where the system is framed in terms of the physical laws that determine the systems behavior, particularly of water and cellular molecules and ions, as well as the mechanical characteristics of the membrane.

### The Standing Gradient Osmotic Flow Model

The model that is most commonly invoked to explain isotonic water transport is the standing gradient osmotic flow (SGOF) model, which was devised by Diamond and Bossert (1967). It proposes that the very small spaces between epithelial cells, the ‘lateral intercellular spaces’ (LISs), are essential for driving water across epithelia. On the apical side the LISs are bounded by tight junctions, while they are open on the basolateral side. The SGOF model hypothesizes that NKAs are situated mostly on the lateral membranes of the epithelia and drive Na^+^ into the LIS so that the osmolarity there is elevated relative to the intracellular and extracellular solutions. The smallness of the LIS facilitates this increase and draws water from the apical side. This leads to a flow of solution through the LIS into the extracellular space surrounding the basolateral surface of the epithelium.

However, several investigators have suggested that it is possible to account for the movement of water by the sodium pump developing very small osmolarity gradients, so small as to not be measurable (Mathias and Wang, 2005; Spring, 1983), without the necessity to invoke the LISs. Mathias and Wong’s model was phenomenological and did not explicitly model the fluxes of ions. Here, we provide a model that includes ionic fluxes. One of the implications of our analysis is that the operation of the NKA can lead to minute differences in osmolarity across a membrane that are nevertheless sufficient to drive water transport. These differences in osmolarity could be so small as to be well below the limits of detection by current techniques.

### Boundary conditions

We will show that a key part to understanding what drives water transport is adopting boundary conditions that match the system. In most epithelial simulations two kinds of boundary conditions have been used: 1) **Open-circuit**, where the transepithelial potential is treated as a variable, and both bath concentrations and pressures are given as fixed boundary conditions (Layton and Edwards, 2014; Weinstein, 1992). To find the voltage in the lumenal compartment the transepithelial fluxes are assumed to be zero. 2) **Short-circuit**, where the voltages in the two compartments are fixed, and current can flow through the epithelium but there is no transport of water, emulating conditions under voltage-clamp.

These two conditions can be misleading as epithelia cannot be accurately represented by electrical circuits because epithelia drive water fluxes and allow ion accumulations. As we will show the adoption of these boundary conditions have hampered the understanding of epithelial transport. To rectify this, we suggest the following boundary conditions. 1) **Natural boundary conditions** (NBCs), where the size of the sink compartment is allowed to vary as well as its voltage. Importantly, transepithelial volume fluxes are allowed. 2) **Voltage-clamp with flow** (VCF), where the transepithelial voltage is clamped, and the transepithelial current is measured. In contrast to the short-circuit condition, we allow fluxes into the sink compartment, and its volume to vary rather than making it infinite.

As mentioned above, most simulations have used infinite baths flanking the epithelium. However, there have been a few exceptions, which have allowed the sink compartment to expand in their models (Dvoriashyna et al., 2018; Gin et al., 2007; Verma et al., 2026).

### Voltage clamp and short circuit current

Ussing and Zerahn (1951) realized that if the solutions on both sides of an epithelium in a chamber were the same, no current would flow through the paracellular pathways at a holding potential of zero, since all ionic reversal potentials will be zero. However, if an electrogenic ion transporter is active, a current will be recorded. In the case of the NKA, the net transmembrane cation flux will be registered. However, there will be no accompanying Cl^-^ current. This is because Cl^-^ ions move into the cathode, compensating for cations moving across the membrane from the cathodal compartment, while Cl^-^ ions flow out of the anode compensating for cations flowing into the compartment (Schultz, 1980). However, *in situ* the fluxes into the sink compartment need to be balanced because of the electroneutrality constraint.

### History of modeling of epithelia

A very useful summary of epithelial transport has been given by Tripathi and Boulpaep (1989), which delves into how epithelial ion and water transport are measured, as well as modeling these processes. The first two models of epithelia that were based on the PLM were published in 1979 (Lew et al., 1979; Weinstein and Stephenson, 1979), and although they both essentially implemented PLMs, neither referenced Tosteson and Hoffman’s paper.

Conway and Boyle (1941) were probably the first to realize the importance of charge constraints on the transport of water and ions in single cells. This turns out to be enormously important in comprehending how water can be driven across membranes and was included by both models (Lew et al., 1979; Weinstein and Stephenson, 1979).

### Paracellular pathways

Tight Junctions (TJs) form a barrier between the apical side of an epithelium and the LIS, with their permeability characteristics varying, depending on the nature of the epithelium. The permeability of a TJ is determined by claudin molecules (23 genes in humans), which are transmembrane proteins that form channel like structures between cells (Citi et al., 2024). Depending on the type of claudin, they can prevent the passage of ions and water, allowing only water to flow, or a mix of ions and water. Claudins form ion channels in the space between epithelial cells and are unusual in that they are not embedded in a membrane and hence do not have a parallel capacitance as do classical membrane ion channels. Although the flux of water through claudins was demonstrated fifteen years ago (Rosenthal et al., 2010), their precise role in epithelial transport has remained uncertain. Our analysis demonstrates that they can play an essential role in ion and water transport.

### Epithelial Pump-Leak Model (ePLM)

We first consider a typical tubular epithelial system as shown in Fig. 1 a). We contend that we can, to a first approximation, reduce the system to Fig. 1b, which has three compartments: an infinite extracellular compartment (e) where the ion concentrations are fixed, a compartment *A* with two membranes, 1 which connects *A* with the extracellular compartment through a set of channels and transporters, and 2 which connects *A* with compartment *B* with a different set of channels and transporters. Compartment *A* represents the cells of the epithelium. Compartment *B* represents, depending on the context, either the lumen or the extracellular space adjacent to the basolateral surface of the epithelium. In addition, *B* is connected by a passage to the extracellular compartment through conductances and water permeability, but no membrane, which represents the paracellular pathway. Each cell and space is represented by a single compartment. The assumption is that concentrations and voltages are uniform showing no spatial gradients. For most single cells this is a reasonable assumption, since on the micron scale, diffusion is rapid (Berg, 1993).

**Fig. 1.**
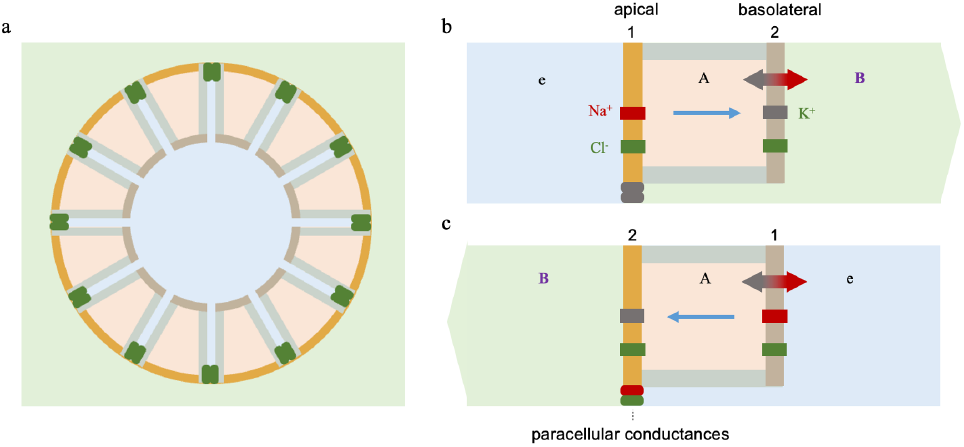
(a) Schematic of a cross-section through a tubular epithelium. (b) KJU model (c) Reverse KJU model. Double headed arrow and rectangles indicate NKA and conductances, respectively.

We have shown (Tarrant et al., 2026) that for certain choices of the conductances, it is possible for both compartment *A* and *B* to be stable. Under these conditions it is possible to have the continuous circulation of Na^+^ and K^+^, but not Cl^-^, at least when there is no active Cl^-^ transport. When the ions pass through channels, they do so with water of hydration. This can also lead to the circulation of water (see below).

Our approach is to search for solutions where the volume of compartment *A* is stable in the steady state, but the volume of compartment *B* increases linearly with time representing transepithelial transport of water and solutes. We will show that for the sort of ion channels and transporters that we consider, the only feasible solution is one with a linear increase in volume. We will refer to this model as the *AB*^*f*^*p*, where *f* indicates that there is continuous flow into *B*. Depending on the context, membrane 1 or 2 can serve as the basolateral membrane, but we always use *B* as the expanding compartment (Fig. 1 b, c).

In this work, we search for solutions where the volume of *B* increases linearly as a function of time. Alternatively, *B* can be considered as a tube with the same diameter as *A*, whose length grows steadily as a function of time representing the transepithelial transport of fluid.

Most models of epithelial transport have modeled compartment *B* as infinite, and in the short-circuit configuration, where the potential in compartment *B* is fixed. In this configuration, the osmolarity in *B* cannot be increased by the action of the NKA. The short-circuit configuration mimics the voltage-clamp experiments that are typically performed in a Ussing chamber. However, we believe that a reliance on these models has obscured the precise mechanism whereby the action of the NKA drives water fluxes. Namely, developing a small gradient in osmolarity and a voltage in compartment *B* different from that in the extracellular compartment. It has also clouded the understanding of the role of the paracellular pathway in water transport.

### Calculating potentials

In the pump-leak framework the calculation of voltages in compartments flows seamlessly from the equations. As Fraser and Huang (2007) have shown, there are two ways to calculate the potential in a compartment: (1) The ‘current-sum’ (CS) method where an expression for the time derivative of the voltage, 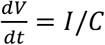, where *I* is the current flow into the compartment, is integrated, and *C* is the capacitance. (2) The ‘charge-difference’ (CD) method where an algebraic equation for *V* is used, *V* = *Q*/*C*, where *Q* is the total charge in the cell. It needs to be emphasized that this method is exact and simple to implement. But oddly it has not been widely used before in the context of epithelial models. In most epithelial models the voltage has been calculated using minimization to find a voltage that makes the transepithelial flux zero, which is cumbersome and unnecessary.

The CD form of the voltage simplifies solution of the set of equations that encapsulates epithelial function, and it is unnecessary as some do, to append an equivalent circuit model to calculate the voltage (Larsen and Sørensen, 2019). In addition, it is easier to include impermeant molecules and their charges when one uses a CD form. Perhaps most significantly the CD form allows one to calculate the voltage in the sink compartment, when it is ‘floating’, that is when both the voltage and volume of the sink can vary. This is not the same as the ‘open-circuit’ which sometimes purported to be the condition *in situ* (Sackin and Palmer, 2013).

### Electroneutrality

Electroneutrality is often applied as a constraint to solve pump-leak equations (PLEs). However, it simply emerges from the interaction of the forces at play in an osmo-electrodiffusion system. If one sets up the system according to physical laws *(Development of the ABp equations*), there is no explicit constraint, however, electroneutrality emerges out of the physics of the situation. Moreover, in the steady state the compartments are only approximately electroneutral. If there were exact electroneutrality, no transmembrane voltages could arise, and the mechanism for action potential generation would be impossible.

If one considers a single compartment PLM, the disparity from electroneutrality increases as the capacitance increases, which allow the charges to spread out and minimize their interactions. The greater the area of the membrane the larger the charge that is needed to generate a given voltage.

It has been claimed that unless electroneutrality is imposed on the system, because of the ion fluxes, it will violate electroneutrality (Latta et al., 1984). However, if the forces and fluxes are correctly specified the compartments move to an approximately electroneutral steady state.

Most models of epithelial transport have excluded consideration of the membrane capacitance, with some exceptions (Dvoriashyna et al., 2018). Because the lipid bilayer behaves like a dielectric, all cells have capacitive elements. The capacitance does not enter the steady state solution. However, inclusion of the capacitance greatly simplifies the calculation of membrane potentials.

## Results

### Development of the ABp equations

In this section we show how to derive the set of equations that encapsulate the behaviour of an *AB*^*f*^ *p* system, from physical laws.

1. The osmotic flux of water is determined by the difference in osmolarity between two compartments. For each membrane, and paracellular pathway, there is an equation of the following form:

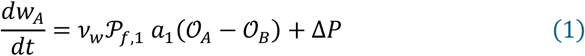

where *w*_*A*_ is the volume of compartment *A, v*_*w*_ is the partial molar volume of water, *P*_*w*,1_ is the osmotic permeability of membrane 1 (Finkelstein, 1987), and *a*_1_ is its area. Δ*P* is the hydrostatic pressure across the membrane. The osmolarity of the *j*^*th*^ compartment is give by:

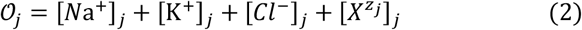

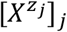 is the concentration of all impermeant species in compartment *j*. We assume that the cell membrane is pliant and can be stretched indefinitely. Equation 1 is often termed Starling’s equation and is widely assumed to hold true in physiology (Blaustein et al., 2019). This equation only holds true for the uncharged molecules and may breakdown when charged molecules are included (Manning, 2026).
2. The voltage within the *j*^*th*^ compartment is proportional to the sum of all charges in that compartment.

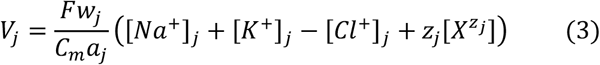

*z*_*j*_ is the average charge on all impermeant molecules in compartment *j* = *A, B*. Notice that in determining *z*_*j*_ one includes all impermeant molecules, even uncharged ones. *C*_*m*_ is the unit membrane capacitance and *a*_*j*_ the area of the *j*^*th*^ membrane.
3. The rates at which ion concentrations change within a compartment are determined by Ohm’s and Fick’s laws, that govern the rates of efflux and influx into that compartment. These fluxes are set by the ion channels and active ion pumps connecting the compartment to those surrounding it. See Equations A6 and A7 in Appendix A.

This type of model includes electrodiffusive fluxes of ions, the osmotic flow of water and changes in compartment volume. We will hence refer to these as osmo-electrodiffusion (OED) equations. In what follows we will simply assume that the membrane is pliant, however one can include the mechanics of the membrane and cytoskeleton if need be (Venkova et al., 2022). The full equations used in this paper are given in Appenix A and the default values of the parameters are given in Table 1.

**Table 1.**
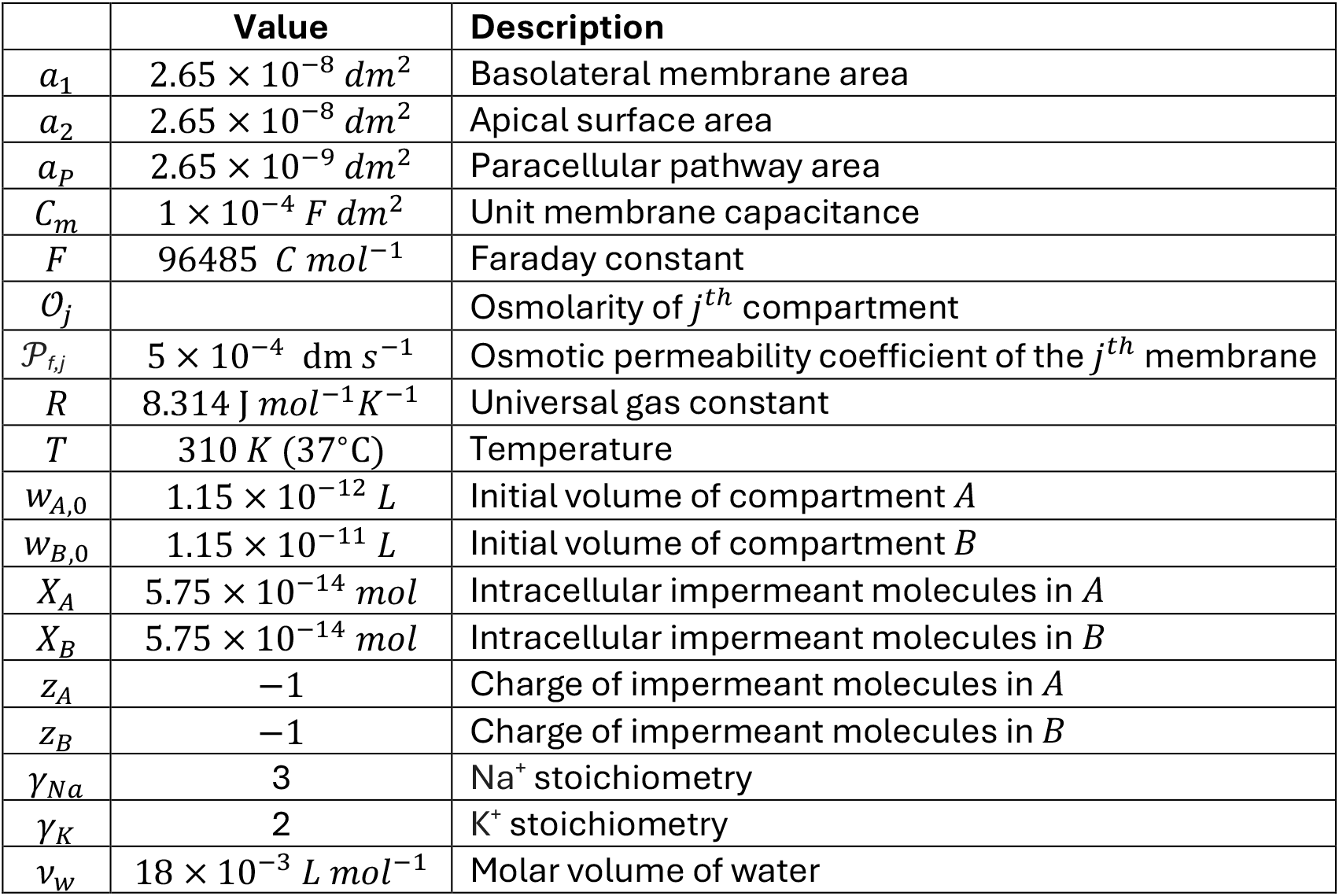
Parameters and variables.

**Table 2.** Composition of the extracellular solution.

|  | Value | Description |
| --- | --- | --- |
| $\mathcal{O}_e$ | 300 mOsm | Extracellular osmolarity |
| $[Y^{ZY}]_e$ | 1 mM | Extracellular impermeant molecule |
| $z_Y$ | -1 | Charge of extracellular impermeant molecule |
| $[Na^+]_e$ | 147 mM | Extracellular Na<sup>+</sup> concentration |
| $[K^+]_e$ | 3 mM | Extracellular K<sup>+</sup> concentration |
| $[Cl^-]_e$ | 149 mM | Extracellular Cl<sup>-</sup> concentration |

**Table 3.** Conductances in S dm^-2^.

| | KJU | rKJU | $\ell$ KJU | Description |
| --- | --- | --- | --- | --- |
| $g_{Na,1}$ | $0.005 a_1$ | $0.005 a_1$ | $0.005 a_1$ | Na<sup>+</sup> on membrane 1 |
| $g_{Na,2}$ | 0 | 0 | $0.0005 a_2$ | Na<sup>+</sup> membrane 2 |
| $g_{Na,p}$ | 0 | $5 a_p$ | 0 | Na<sup>+</sup> on para. pathway |
| $g_{K,1}$ | 0 | 0 | $0.001 a_1$ | K<sup>+</sup> on membrane 1 |
| $g_{K,2}$ | $0.03 a_2$ | $0.03 a_2$ | $0.03 a_2$ | K<sup>+</sup> on membrane 2 |
| $g_{K,p}$ | $5 a_p$ | 0 | $5 a_p$ | K<sup>+</sup> on paracellular pathway |
| $g_{Cl,1}$ | $0.1 a_1$ | $0.1 a_1$ | $0.1 a_1$ | Cl<sup>-</sup> on membrane 1 |
| $g_{Cl,2}$ | $0.1 a_2$ | $0.1 a_2$ | $0.1 a_2$ | Cl<sup>-</sup> on membrane 2 |
| $g_{Cl,p}$ | 0 | $5 a_p$ | $5 a_p$ | Cl<sup>-</sup> on paracellular pathway |

$\ell$ KJU – loose KJU

Epithelial operate in a dynamic steady state, where over extended periods of time, intracellular ion concentrations and cellular volumes remain fairly constant. The objectives of OED models are to replicate this steady state. The goal is to find the set of ion channels and pumps that can stabilize an epithelium, and capture the steady state ion concentrations and fluxes. The stability of such a system is by no means guaranteed, since some parameter combinations lead to a continuous increase in cell volume or to the volume collapsing (Mori, 2012). What we ultimately would like to do is to find the regions of parameter space that support stable operation of the epithelium.

There have been a good number of simulations using this approach, however what is new here is the introduction of Equation (2), which considerably simplifies solution of the equation. This may seem like a small change, however, it allows one to find the transient solutions to OED equations and include more natural boundary conditions. Namely, it is simple to include cases where the lumenal space is finite and both these spaces and the cells can change volume. Moreover, we have been able to find analytical solutions to the *AB*^*f*^*p* system (Appendix B and C). In what follows we consider an A*Bp* model with the NKA only on the basolateral membrane, and Cl^-^ conductances on the apical and basolateral membranes, while various combinations of Na^+^ and K^+^ conductances are considered on the two membranes (Fig. 1 b, c). In addition, we allow for different combinations of paracellular conductances. In appendix B we derive analytical expressions for the *AB*^*f*^*p* system where the volume of *B* grows linearly with time. We show that depending on the balance of Na^+^ and K^+^ conductances the water can be transported from the basolateral to the apical side (absorptive mode, Fig. 1 b) or in the opposite direction (secretory mode, Fig. 1 c).

One of our central findings is that the constraints on flow are very different for *A* and *B*. For *A*, because ions can move transepithelialy, there can be arbitrary flows of ions through it, as long as the inflow of a given ion matches that of its outflow. However, the different ion flows do not need to balance. In contrast, for compartment *B*, because it grows in volume, it has to have a net influx of ions into it. However, to preserve rough electroneutrality the influx of positive and negative ions has to balance. To show this we have developed what we term a *flow diagram*, where the heights of the rectangles represents the magnitude of the ion flows. A hypothetical example is shown in Fig. 2, where only K^+^ and Cl^-^ ions flow into and out of *A* and flow into *B*. The flow into and out of *A* are not balanced. To balance the flow into *B*, Na^+^ and Cl^-^ flow through their paracellular pathways, making the total flow of positive charges into *B* equal to that of the negative charges

**Fig. 2.**
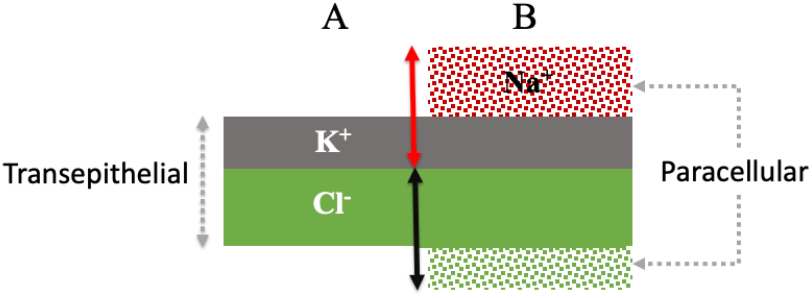
Flow diagram for a hypothetical *AB*^*f*^*p* system. The flow for all ions is from left to right. The red and black arrows show that the flow of cations into *B* equals that of the anions. The paracellular flow is indicated by stippled rectangles, while the transepithelial flow is indicated by solid ones.

### Koefoed-Johnsen-Ussing Model

We first consider the classical Koefoed-Johnsen-Ussing (KJU) (1958) model shown in Fig. 1 b, where the NKA is on the basolateral membrane, which has a high K^+^ conductance and is in contact with compartment *B*. While the apical membrane is permeable to Na^+^ but not to K^+^, both the apical and basolateral membranes are permeable to Cl^-^. Compartment *B* is connected through a paracellular pathway to the extracellular space, although this was not considered in the original KJU model, nor did they include impermeant molecules, or develop a mathematical model.

For the KJU model, for a range of conductances, steady states are possible where *A* is stable and *B* grows at a constant rate, reflecting the continuous flow of solution from the luminal side through the apical membrane and then through the basolateral membrane into the extracellular space (Fig. 3 a). The approach to steady state is shown in Fig. 3 a with continuous linear growth of compartment *B*.

**Fig. 3.**
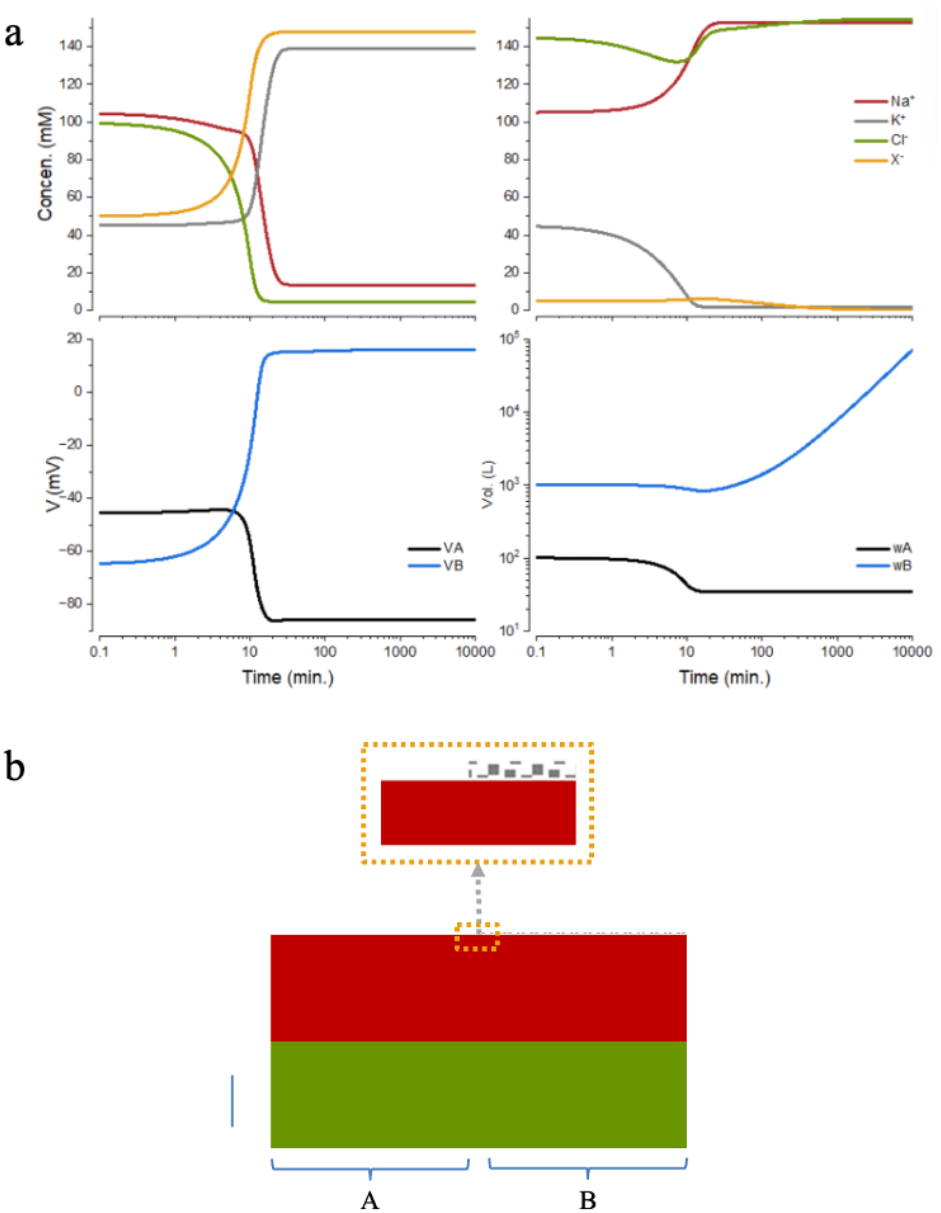
a) Approach to steady state of KJU [010]. b) The flow diagram for the KJU. The height of the color band represents the magnitude of the flow. The inset shows an enlarged version of the section in the box. Stippled field is flow through the paracellular pathway p = 2.5 x 10^-4^ A dm^-2^.

What becomes clear for the KJU model is that paracellular pathways are necessary for the flow and the stability of *A*. For each ion, the flow into and out of *A* must be balanced for *A* to be stable, but these flows do not need to balance one another. On the other hand, because the water flow into *B* must be continuous to ensure epithelial transport, the steady state concentrations in *B* should be stable. Otherwise, it would be impossible to maintain a stable *A*. Moreover, the flow of positive and negative charges into *B* must be balanced to ensure that the solution remains electroneutral. This is not possible in the case where there is no paracellular pathway (see a proof in Appendix B).

In the case of the KJU, it works best if there is only a K^+^ flux through the paracellular pathway, which can be seen in the flow diagram in Fig. 3 b. The flux through this pathway is very small but if it is absent the water transport system breaks down.

To designate whether a paracellular pathway is open or closed to an ion flow, we use 1 or 0 respectively, writing it as [Na^+^ K^+^ Cl^-^]. For the KJU model there are four combinations that permit stable flow: [011], [010], [111], and [110], in the order of decreasing flow rate.

Two combinations that lead to no flow but stable *A* and *B* are [100] and [101]; we will refer to this as a ‘jammed’ system. Lastly two combinations where *A* is unstable are [001] and [000].

A key question is what drives the vectorial water transport. It is clear in this model that differences in osmolarity can be very small and are maintained by the operation of the NKA (*O*_*A*_ = 304.7 mOsm, *O*_*B*_ =309.4 mOsm). These differences from the extracellular osmolarity (300 mOsm) may be too small to be measured, but they are sufficient to generate fluxes of the magnitude that have been detected. Water fluxes of ∼ 2.3 *μL cm*^−2^*h*^−1^ have been measured across frog skin (Steinbach, 1967). For the KJU model in Fig. 3 the steady state water flux is 1.9 *μL cm*^−2^*h*^−1^. Note that unlike the SGOF model it is not necessary to invoke a small space like the LIS to support water fluxes.

We consider the case shown in Fig. 3 with a higher *P*_*f*_ (5 x 10^-3^ dm s^-1^). In the steady state *O*_*A*_= 300.484 mOsm, *O*_*A*_ = 300.967 mOsm, and the water flow rate is 1.38 fL s^-1^, which corresponds to 76.7 f mole s^-1^ for water molecules (*w*_*A*_ = 395 f L). The total flow rate of all ions into compartment *B* is 0.42 f mole s^-1^. So, each ion that moves into *B* carries 184 water molecules by osmosis.

It is clear in our model that the flux of ions driven by the NKA is impelling the water flow; water simply follows the osmotic gradient. However, the model does not include the direct interaction of ions and water. It is known that for K^+^ to pass through voltage-gated K^+^-channels, the water of hydration is removed as it passes through the channel (Pfeffermann et al., 2021). In contrast, Na^+^ ions moving through voltage gated Na^+^ channels hold on to some of the water of hydration (Hille, 1971). If this holds true for leak channels, this could increase the transepithelial water transport. If water is carried through channels with ions, that is what is known as ‘electroosmosis’ (Finkelstein, 1987).

### Loose KJU

It is unlikely that any of the conductances are zero, so we relax this requirement to get what we call the ‘loose KJU’. If we allow *g*_01/_ and *g*_*K*1_ to be finite, the system can still transport water from the apical to basolateral side (Fig. *4* a). Note that K^+^ flows from the basolateral side to the apical side through *A*, while all other ions and water flow in the opposite direction. In the case of the loose KJU all combinations of paracellular conductances are stable, but they do still have an influence on the rate at which water is transported. The highest rate is with [011] and falls off in the order [010], [001], [000], [111], [101], [110], and [100]. The rate of water transport is strongly influenced by *gna*_1_, less so by the other conductances.

### Reverse KJU

The direction of water transport in the KJU can be reversed by exchanging the values of the Na^+^ and K^+^ conductances on the apical to those of the basolateral side (Fig. 1c) and vice versa. Now water flows from basolateral to the apical side of the epithelium. We will call this the reverse KJU (rKJU). Notice here that most of the ionic flux occurs through the paracellular pathway (Fig. 4 b). As with the KJU, a loose rKJU functions as well, but we will not consider this here.

**Fig. 4.**
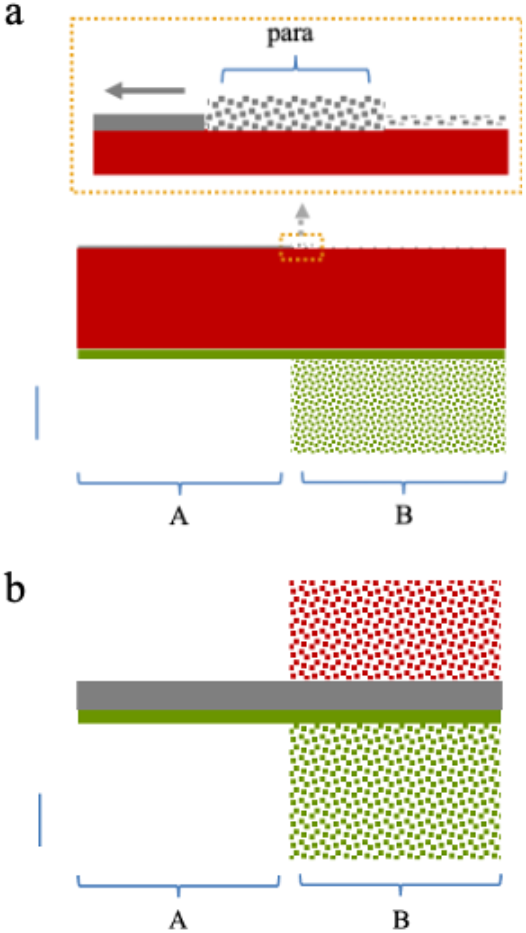
Flow diagrams of (a) loose KJU ([011], p = 2.5 x 10^-4^ A dm^-2^ ) and (b) reverse KJU ([1 0 1], p = 1 x 10^-4^ A dm^-2^). For (a) note that the flow of K^+^ through *A* is from right to left.

### Analytical solutions

The analytical solutions to the *AB*^*f*^*p* system, which are derived in Appendix B, facilitate a global view of the systems’ behavior. We have confirmed that the analytical solutions match those calculated by numerical integration (data not shown).

Water transport is very dependent on the osmotic permeabilities (*P*_*f*_) of the three pathways; the apical, basolateral, and paracellular routes. We will assume that they have equal permeability coefficients. The *P*_*f*_ of lipid bilayers has been found to be in the range of 1.6 - 8.3 x 10^-4^ dm s^-1^. We will assume *P*_*f*_ = 5 x 10^-4^dm s^-1^, and consider what happens when it is increased by an order of magnitude. In Fig. *5* we have plotted the concentration of the ions in *A* and *B*, osmolarities, voltages, the rate of water transport, and the balance of cation and anion flows into *B*, as functions of the pump rate, what we have previously termed a *cP* plot (Kay, 2017). As the rate of the NKA is increased the concentration of K^+^ increases intracellularly, and the rate of water transport rises, driven by the differences in osmolarity between the compartments (*B*>*A*>*e*), which can be very small. Note that the rise in water transport rate (*m*_*B*_) is almost linear and the rise persists even when the concentration of K^+^ in *B* asymptotes. At low pump rates the osmolarity of the solution transported into *B* is only very slightly higher than that of the extracellular solution until the pump rate becomes so high its osmolarity exceeds that of the extracellular solution. As *P*_*f*_ is decreased the action of the NKA can sustain progressively higher differences in osmolarity between the compartments. With the highest *P*_*f*_ used here, at high NKA pump rates it is possible to sustain large differences in osmolarity between the compartments. The bottom panels of the figure demonstrate that the influx of cations and anions into *B* are balanced throughout the range of pump rates considered.

**Fig. 5.**
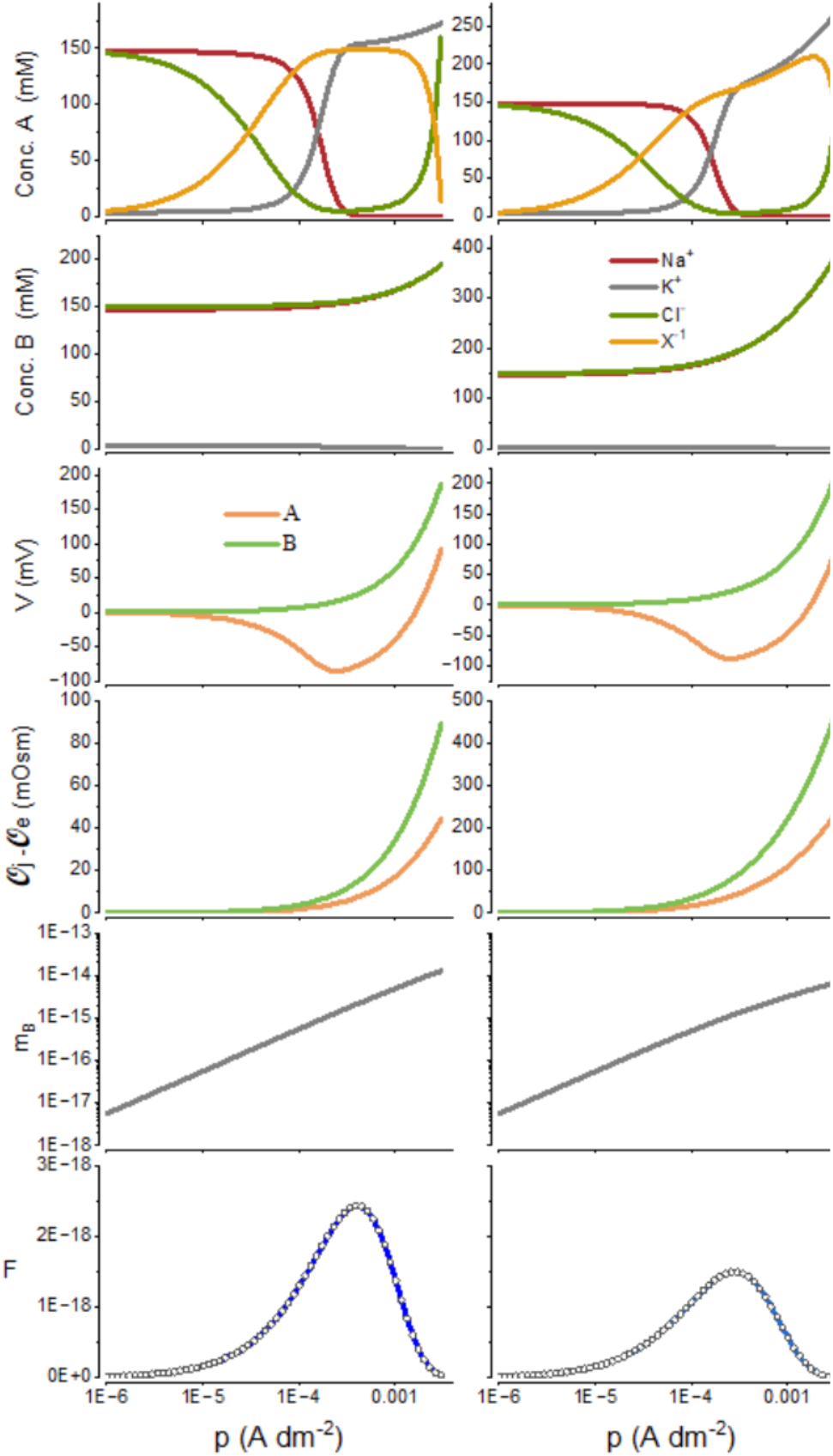
cP plots for the KJU model derived from analytical solutions of the *AB*^*f*^*p* equations. Left column *P*f = 5 x 10^-4^, right column *P*f = 5 x 10^-5^ dm s^-1^. The fourth row shows the difference between the osmolarity in a compartment and the extracellular osmolarity. *m*_*B*_ is the rate of water transport into *B*. The bottom panel shows the difference between the transepithelial Cl- and Na^+^ flow into *B* (solid line) and the paracellular K^+^ flow into *B* (circles).

The analytical solutions that we derived have an upper bound for the pump rate, *p*_*max*_, above which the system becomes unstable. *p*_*max*_ can be computed from the analytical solution to the PLEs (see Appendices B and C). We noticed that the apparent *p*_*max*_*s* when numerical methods were used to solve the equations, were lower than that of the analytical solution. For the KJU model with the same parameters as shown in Fig. 3, the numerical *p*_*max*_ is ∼ 1.68 × 10^−3^ *A dm*^−2^, while it is ∼ 3 × 10^−3^*A dm*^−2^ for the analytical solution. However, because *p* = *p*_*max*_ corresponds to a singular point of the system, these steady states may not be numerically accessible or reliably resolved. Thus, failure to obtain a numerical solution at *p* = *p*_*max*_ should not be interpreted as evidence that a steady state does not exist.

Unlike the KJU model the rKJU model with high water permeabilities produces a transportate that differs from the extracellular solution even at very low pump rates (Fig. *6*).

**Fig. 6.**
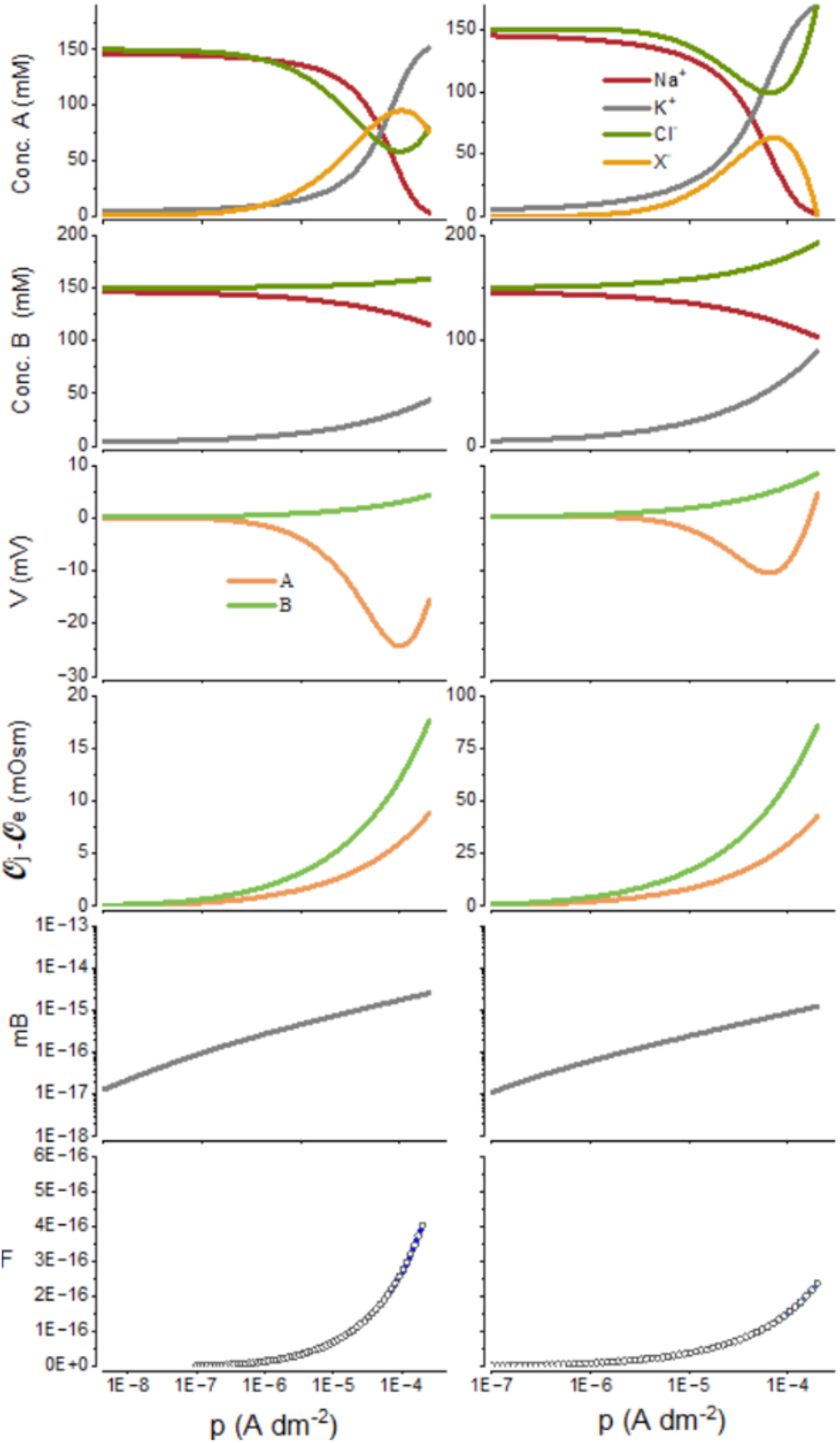
*cP* plots for the rKJU model derived from analytical solutions of the *AB*^*f*^*p* equations for *P*f = 5 x 10^-4^ (left column) and *P*f = 5 x 10^-5^ dm s^-1^ (right column). The fourth row shows the difference between the osmolarity in a compartment and the extracellular osmolarity. *m*_*B*_ is the rate of water transport into *B*. The bottom panel shows the flow of Na^+^ and K^+^ into *B* (solid line), and the flow of Cl-into *B* (circles).

To determine likely pump rates we combine the estimated turnover rate of the NKA (∼ 200 cycles per second) (Lüpfert et al., 2001) with its estimated densities. The highest densities of the NKA are found in renal tubules (∼6,500 per *μm*^2^) (Deguchi et al., 1977; El Mernissi and Doucet, 1984), which would generate a *p*∽2.1 × 10^−3^ *A dm*^−2^. If the NKAs were packed at the maximum possible density, which is about 12,000 per *μm*^2^, then *p*∽3.9 × 10^−3^ *A dm*^−2^.

### Although Cl^-^ is not actively transported, it is not at equilibrium

In a single compartment PLM model, if Cl^-^ is not actively pumped, the chloride distributes passively according to the membrane potential (i.e., *V* = *E*_*cl*_) (Jakobsson, 1980). However, in the *AB*^*f*^*p* this cannot occur if the potentials across the apical and basolateral membranes are different. In this model these potentials can be calculated from the expression

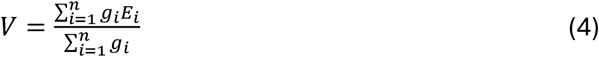

which is accurate for low pump rates. In the *AB*^*f*^*p* model one of the effects of operating the NKA is to pull Cl^-^ out of equilibrium even though it is not driven directly. If the membrane potentials across the apical and basolateral membranes are different, the Cl^-^ concentration in *A* will be simultaneously drawn towards two different equilibria, and hence the concentration will settle into a non-equilibrium steady state.

### Effect of extracellular osmolarity

We have found that the ePLM can operate over a very wide range of extracellular osmolarities (*O*_*e*_). For the KJU, as *O*_*e*_ increases the rate that water is transported (∝ *O*_*B*_ − *O*_*A*_) declines monotonically, with a fixed NKA pump rate (Fig. 7). It does so because as *O*_*e*_ increases it requires more energy to develop the concentration that is required to move the water. It is noteworthy that at low extracellular osmolarities it is possible to establish osmolarities in *A* and *B* that greatly exceed that in the extracellular compartment (Fig. 7, top panel).

**Fig. 7.**
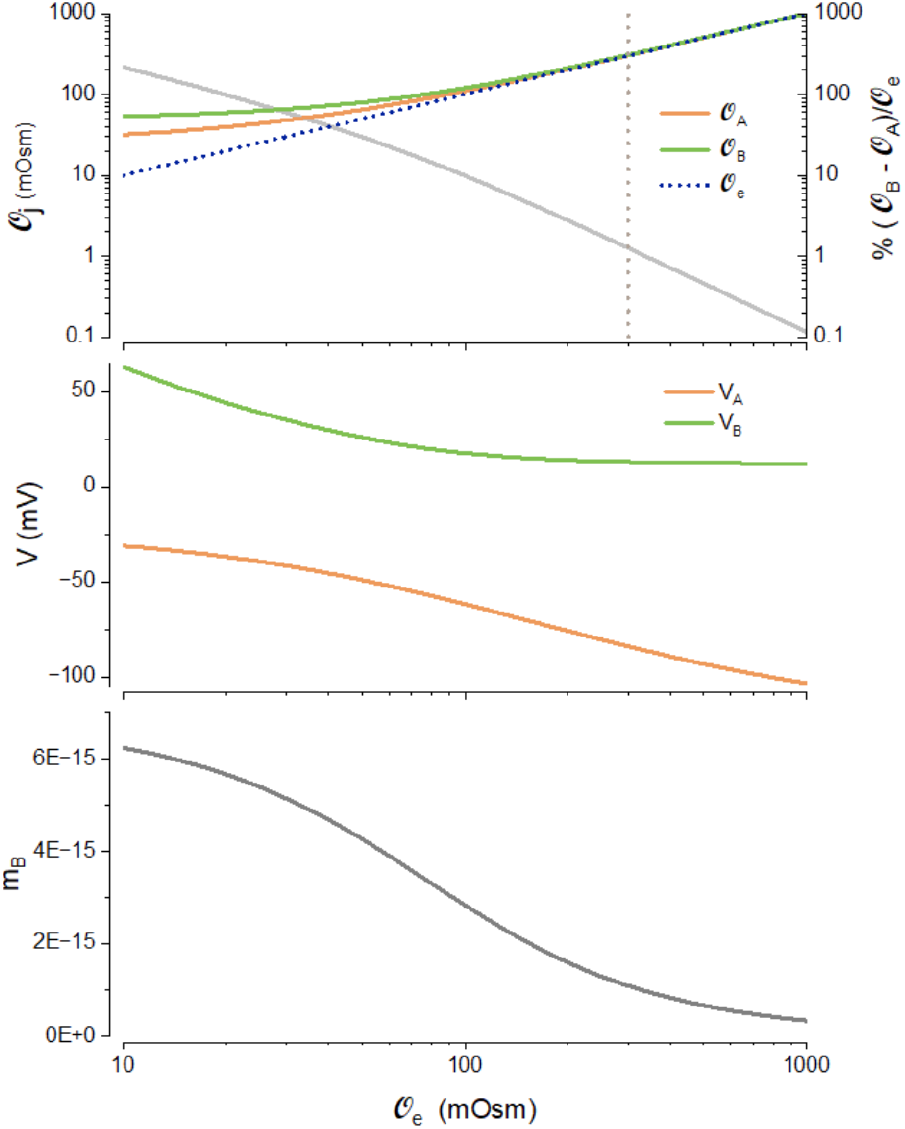
Effect of varying the extracellular osmolarity on the KJU (analytical solutions). *P*f = 5 x 10^-4^ dm s^-1^, p = 2 x 10^-4^ A dm^-2^. The vertical dotted line is at 300 mOsm.

To explore how the PLM operates under conditions where there is an asymmetry between the two flanking compartments, we assume the size of *B* be infinite and the concentrations in this compartment do not vary (Fig. 8) and explore what happens in voltage clamp. The osmolarity of *B* can change from 10 to 1000 mOsm and the system remains stable. When the osmolarity is balanced, there is only a net transport of Na^+^ from the extracellular space to *B*, with no water transport. While, if the osmolarity of *B* is less than 300 mOsm, in addition to Na^+^ transport, Cl^-^ is transported from the extracellular compartment to *B*, while water moves in the opposite direction. In contrast, if the osmolarity of *B* is greater than 300 mOsm, Cl^-^ is transported from *B* to the extracellular compartment, while water moves in the opposite direction. In both the last two cases Na^+^ is transported from e to *B*. The system moves to a steady state where the movement of water across the two membranes is balanced, so that the volume of *A* is constant, by converging to an intracellular osmolarity that makes this possible.

**Fig. 8.**
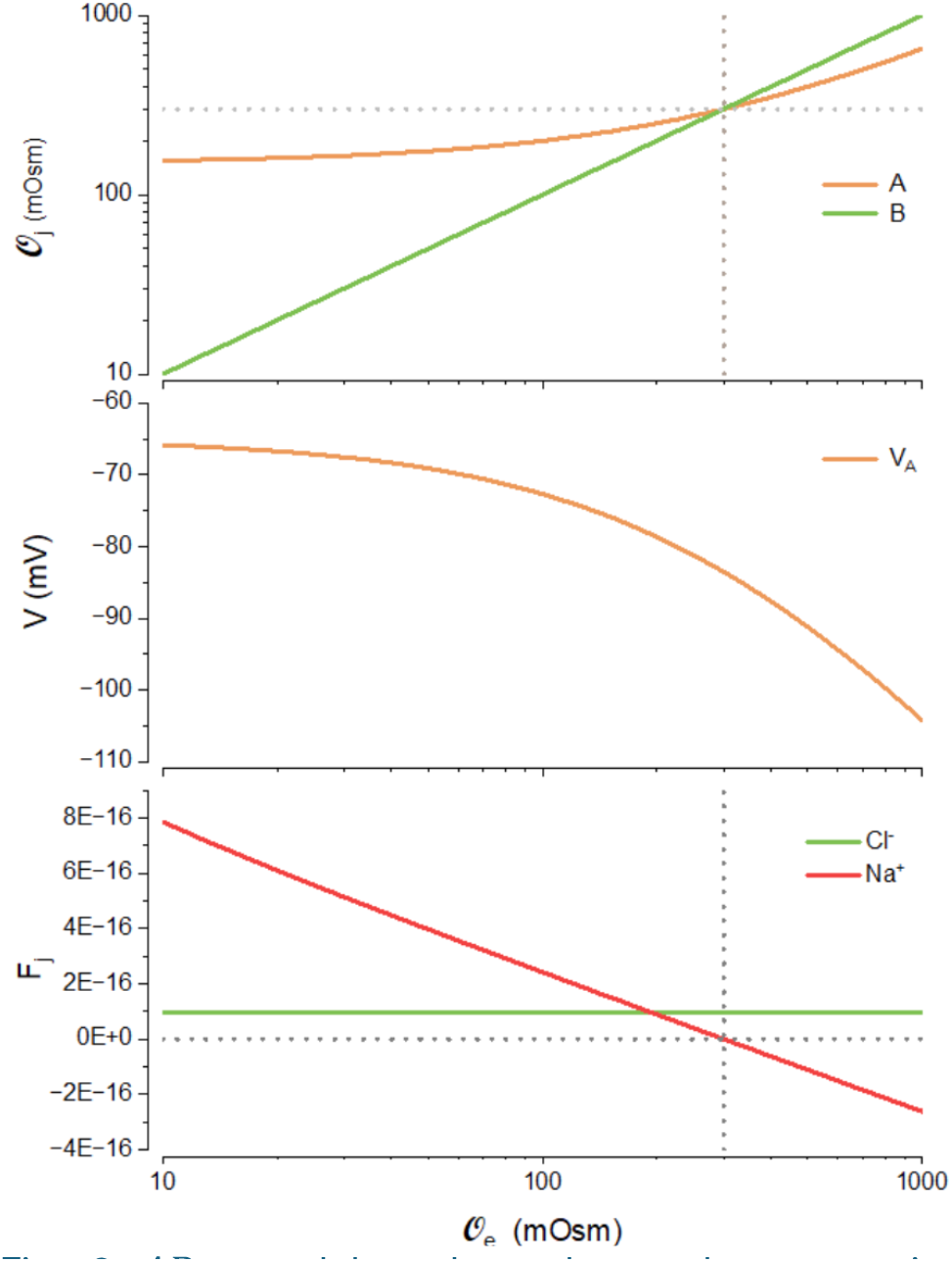
*ABp* model under voltage clamp, varying osmolarity of numerical solution in *B*. Lower panel shows the transepithelial Na+ and Cl^-^ flows. *P*f = 5 x 10^-4^ dm s^-1^, p = 2 x 10^-4^ A dm^-2^

## Discussion

We have shown how the PLM of Tosteson and Hoffman (1960) can be extended to account for water and ion fluxes across an epithelium. Our analysis demonstrates how the action of the NKA can establish osmotic gradients that drive water fluxes. We have shown that it is not necessary to include morphological elaborations like the lateral interstitial spaces to account for water movement. A key part of our analysis was in demonstrating the importance of finding the appropriate boundary conditions to reveal the natural behavior of epithelial.

Tight junctions have long been known to establish patent epithelial barriers between the flanking compartments. Only more recently it has been shown that the ionic and water permeability of tight junctions are controlled by the expression of specific claudin molecules. Although it has long been known that currents can flow through tight junctions, their roles in transport have remained enigmatic. Here we have shown for the first time that tight junctions could serve to balance the flux of cations and anions into the transportate. Moreover, the precise combinations of conductances expressed will have a strong influence on ion and water transport.

Much of theoretical biology relies on the numerical solution of sets of differential equations. This allows one to explore the dynamics of physiological systems, however one is required to inch one’s way through what is sometimes a very large dimensional parameter space. This has the disadvantage that one might miss features unless one chances upon them. Here we have derived analytical solutions to the *ABp* equations that facilitate global insights into the systems’ behavior. Without the analytical solution we would have missed the fact that the ePLM can operate at very low extracellular osmolarities, which has very important implications for animal osmoregulation (Fig. 7 and below).

Although equivalent circuit models can explain some aspect of epithelial physiology (Leaf et al., 1981), we believe their adoption has hobbled our understanding of water transport. It has done so because equivalent circuits can neither, at least in their standard form, include the accumulation and redistribution of ions, nor volume changes that result from water fluxes.

The CD method provides an excellent way of modeling the transport dynamics of epithelia. In most epithelial simulations minimization procedures have been employed to solve the the open-circuit case. Here we have shown that it is considerably simpler to use the CD method to arrive at a solution, without applying unrealistic boundary conditions. Another advantage of the CD method is that it allows for the solution of the transient behavior of the system. Central to the application of the CD method are the cells’ capacitance, which have typically been omitted from epithelial models.

### Electroosmosis

There has been some dispute as to whether electroosmosis could play a role in water transport (Fischbarg, 2010; Hill, 2008). Part of the problem is trying to disentangle it from osmosis and we believe that the ePLM can be of value in clarifying this question.

In the ePLM used here, water does not move through ion channels. However, if Na^+^ ions move through their channels with water of hydration, this will add to the net flux of water, since there is a net polarized flux of Na^+^. In this case water is transported both by osmosis and electroosmosis.

It is difficult to conceive of how one could measure the percentage of water that is carried by electroosmosis. It would require that one be able to turn off the water passage through Na^+^ channels, without affecting the ion passage. Na^+^ channels certainly can be blocked, however that would prevent the operation of the PLM. It is very unlikely that one could mutate Na^+^ channels to eliminate water passage, since the movement of hydrated Na^+^ ions forms the very basis of the monovalent selectivity of the channel (Hille, 1971). Having a model like the ePLM throws into sharp relief precisely how difficult it is to separate electroosmosis from osmosis.

Epithelial physiologists talk about solvent drag, where the movement of water through say an aquaporin or claudin channel could drag ions with it. This has been postulated to occur in the proximal tubule of the nephron (Weinstein, 1992).

It is certainly possible that other variables such as pressure, or morphological complexities may be important. However, if it is possible to account for isotonic water transport with the simple arrangement of cells and extracellular spaces that we have presented, their addition is unnecessary.

### Physiological implications of the ePLM

The PLM has primarily been used to simulate environments where the extracellular osmolarity is around 300 mOsm. Here we have shown that the model can operate at low extracellular osmolarities, which are particularly important for animals that live in freshwater lakes. We have shown that for *O*_*e*_∽ 300 *mOsm*, it is possible to drive the osmolarity in *A* and *B* to slightly higher osmolarities, with biologically realistic NKA pump rates. However, with these same pump rates, at low *O*_*e*_, it is possible to elevate the osmolarities of *A* and *B* more substantially (Fig. 7). This is important because it offers a possible key to understanding how animals transitioned from salt water to fresh water and vice versa (Whittamore, 2012). It shows that an ePLM operating with a realistic energy budget, dwelling in a water with osmolarity of ∼ 10 mOsm, can elevate intracellular osmolarities to 200-300 mOsm. This is crucial because there are no cells in the animal kingdom, as far as we know, that operate below this range (Burton, 1983).

It is a long-held assumption in the field of osmoregulation that the intracellular osmolarity closely matches that of the extracellular osmolarity (Guyton and Hall, 2025; Strange, 2004). Here we have shown that it is quite possible to have a sustained difference between the intracellular and extracellular osmolarity without developing a hydrostatic pressure (Fig. 7). No pressure builds up in *B* because the compartment can expand continuously if there is no impediment to flow. In the steady state no pressure develops in *A* since its volume remains constant.

It is worth noting that a sustained difference between the intracellular and extracellular osmolarities would be unlikely to occur in non-epithelial animal cell, like a blood cell, in a large extracellular solution. To see why this is so, consider a cell where *O*_*e*_ is greater than *O*_*A*_, water would move out of *A* by osmosis. However, to stabilize the volume of *A*, water would have to flow back into *A* to maintain its volume. But the difference in osmolarities would prevent this from occurring. If there were active water pumps this might make it possible, but there is little evidence for their existence (Boron and Boulpaep, 2016).

It is interesting to contrast the evolution of our understanding of cellular excitability and epithelial transport in physiology. The theoretical basis of cellular excitability has been well developed and forms a core of our understanding of how the brain operates (Jack et al., 1971; Sterrat et al., 2011). It is puzzling that this has only occurred to a limited extent in the case of epithelia, where the same principles operate, save for the introduction of water fluxes and hydrostatic pressures. It is even more puzzling given that one of the central mechanisms in kidney function, the counter current multiplier, required a well-developed theory to demonstrate its plausibility (Kuhn and Ryffel, 1942).

Interpretations of experiments inevitably rely on some theory whether it be explicit or implicit. However, if the theory is incorrectly or incompletely formulated, interpretations based on it, collapse. For example, it has been claimed in the case of the choroid plexus that it is not possible without a significant osmotic gradient to drive the formation of CSF (Macaulay, 2021). Our results suggest that this may not be the case.

It is estimated that there are about 150 distinct epithelia in mammals (Caceres et al., 2017). Since there are no documented cases of epithelia without the NKA throughout the animal kingdom, it seems likely that the general principles that we have proposed here should apply to all epithelia.

## Materials and Methods

The coupled differential–algebraic pump leak equations were solved in MATLAB (MathWorks, Natick, MA, version R2025a) using the stiff differential equation solver `ode15s`, with relative and absolute tolerances of 5×10^-6^ and 1×10^-16^, respectively. To improve numerical stability and prevent the solver from generating nonphysical negative values of ion concentrations or compartment volumes during integration, the positive state variables were reformulated in logarithmic form. Specifically, for each physical state variable *X*_*i*_ (including Na^+^, K^+^, and Cl^−^ concentrations and compartment volumes), the numerical state variable was defined as *U*_*i*_ = ln *X*_*i*_, such that 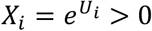 throughout the integration. Accordingly, the original equations *dX*_*i*_/*dt* = *f*_*i*_(*X*) were transformed according to *U*_*i*_/*dt* = (1/*X*_*i*_)(*U*_*i*_/*dt*) = *f*_*i*_ (*X*)/*X*_*i*_, `ode15s` was applied to the resulting log-state equations. This transformation enforces positivity of the physical state variables while retaining the dynamics of the original equations. Graphs were prepared using Origin (OriginLabs, Northampton, MA).

Steady-state solutions theoretically exist at *p* = *p*_*max*_. However, because *p* = *p*_*max*_ corresponds to a singular point of the system, these steady states may not be numerically accessible or reliably resolved. Thus, failure to obtain a numerical solution at *p* = *p*_*max*_ should not be interpreted as evidence that a steady state does not exist.

## Abbreviations

CD: charge difference
ePLM: epithelial pump leak model
KJU: Koefoed-Johnson-Ussing
LIS: Lateral interstitial spaces
NKA: Na^+^ /K^+^ ATPase
PLE: pump leak equations
PLM: pump leak model
rKJU: reverse Koefoed-Johnson-Ussing
TJ: tight junction

## Acknowledgments

We thank Rodney J Douglas for helpful discussions. We used ChatGPT (OpenAI), Clause (Anthropic), and Google Scholar Labs to help search for prior work on the problem addressed in this paper. This work was supported by the National Science Foundation, grant 2037828.

## Author contributions

A.R.K. and Z.A. designed research; A.R.K. and Z.A. analyzed data; Z.A. derived the equations; ARK: prepared the figures. A.R.K. wrote the paper with input from Z.A.

