## Appendix for "A pump-leak model of epithelia reveals the roles of paracellular pathways in water transport & the importance of boundary conditions"

In what follows we present the pump leak equations used in this paper and derive the steady states of KJU and rKJU systems analytically.

### A. ePLM equations

The extracellular and intracellular osmolarities are respectively:

$$\mathcal{O}_e = \left[\mathrm{Na}^+\right]_e + \left[\mathrm{K}^+\right]_e + \left[\mathrm{Cl}^-\right]_e + [\mathrm{Y}^{z_Y}]_e \tag{A1a}$$
$$\mathcal{O}_j = \left[\mathrm{Na}^+\right]_j + \left[\mathrm{K}^+\right]_j + \left[\mathrm{Cl}^-\right]_j + [\mathrm{X}^{z_j}]_j \tag{A1b}$$

All osmotic coefficients are assumed to be unity.

The flow of water is determined by the differences in osmolarity between the compartments through Starling's equation:

$$\frac{dw_A}{dt} = \underbrace{v_1\left(\mathcal{O}_A - \mathcal{O}_e\right)}_{\text{baso. mem.}} + \underbrace{v_2\left(\mathcal{O}_A - \mathcal{O}_B\right)}_{\text{ap. sur.}} \tag{A2a}$$
$$\frac{dw_B}{dt} = \underbrace{v_P\left(\mathcal{O}_B - \mathcal{O}_e\right)}_{\text{para. pathway}} - \underbrace{v_2\left(\mathcal{O}_A - \mathcal{O}_B\right)}_{\text{ap. sur.}} \tag{A2b}$$

Where

$$v_j = v_w \mathcal{P}_{f,j} a_j$$

The charge in the $\mathrm{j}^{\mathrm{th}}$ compartment and the extracellular compartments are:

$$Q_j = \left[\mathrm{Na}^+\right]_j + \left[\mathrm{K}^+\right]_j - \left[\mathrm{Cl}^-\right]_j + z_j\,[\mathrm{X}^{z_j}]_j$$

We apply the electroneutrality constraint to the extracellular compartment:

$$\left[\mathrm{Na}^+\right]_e + \left[\mathrm{K}^+\right]_e - \left[\mathrm{Cl}^-\right]_e + z_Y\,[\mathrm{Y}^{z_Y}]_e = 0$$

The voltage in the $\mathrm{j}^{\mathrm{th}}$ compartment is given by:

$$V_j = \frac{F w_j}{C_m a_j}\left(\left[\mathrm{Na}^+\right]_j + \left[\mathrm{K}^+\right]_j - \left[\mathrm{Cl}^-\right]_j + z_j\,[\mathrm{X}^{z_j}]_j\right) \tag{A3}$$

The flow into each compartment is determined by the passive flow of ions, which follows Ohm's law and the active transport of ions. The equations governing the concentrations of Na$^+$, K$^+$, and Cl$^-$ in compartment $A$ are given by the following equations.

$$F\frac{d\left(w_A[Na^+]_A\right)}{dt} = -\underbrace{g_{Na,1}\left(V_A - E_{Na,A}\right) + \mathbf{p_{Na,1}}} + \underbrace{\mathbf{p_{Na,2}} + d_{Na}}, \tag{A4a}$$

$$F\frac{d(w_A[K^+]_A)}{dt} = -\underbrace{g_{K,1}\left(V_A - E_{K,A}\right) + \mathbf{p_{K,1}}} + \underbrace{\mathbf{p_{K,2}} + d_K}, \tag{A4b}$$

$$F\frac{d(w_A[Cl^-]_A)}{dt} = \underbrace{g_{Cl,1}\left(V_A - E_{Cl,A}\right)}_{\text{ion total flux across basolateral}} + \underbrace{d_{Cl}}_{\text{ion total flux across apical}}. \tag{A4c}$$

Similarly, the equations for the ionic concentrations in compartment $B$ are

$$F\frac{d\left(w_B[Na^+]_B\right)}{dt} = -\underbrace{g_{Na,P}\left(V_B - E_{Na,B}\right)} - \underbrace{\mathbf{p_{Na,2}} - d_{Na}}, \tag{A5a}$$

$$F\frac{d(w_B[K^+]_B)}{dt} = -\underbrace{g_{K,P}\left(V_B - E_{K,B}\right)} - \underbrace{\mathbf{p_{K,2}} - d_K}, \tag{A5b}$$

$$F\frac{d(w_B[Cl^-]_B)}{dt} = \underbrace{g_{Cl,P}\left(V_B - E_{Cl,B}\right)}_{\text{ion total flux across paracellular}} - \underbrace{d_{Cl}}_{\text{ion total flux across apical}}. \tag{A5c}$$

The term $d_{\text{ion}}$ is the flux across surface 2

$$d_{\text{ion}} = -z_{\text{ion}} g_{\text{ion},2}\left(\left(V_A - E_{\text{ion},A}\right) - \left(V_B - E_{\text{ion},B}\right)\right), \tag{A6}$$

the Nernst potentials are

$$E_{\text{ion},j} = \frac{RT}{z_{\text{ion}}F}\ln\left(\frac{[\text{ion}]_e}{[\text{ion}]_j}\right) \tag{A7}$$

and the active transport is denoted by $\mathbf{p}_{\text{ion},1}$ and $\mathbf{p}_{\text{ion},1}$.

The extracellular osmolarity, $[\mathrm{K}^+]_e$ and $[Y^{z_Y}]_e$ are fixed. The concentrations of Cl$^-$and Na$^+$ then follow from the electroneutrality constraint:

$$\left[\mathrm{Cl}^-\right]_e = 0.5\left(\mathcal{O}_e + (z_Y - 1)\left[Y^{z_Y}\right]_e\right) \tag{A8a}$$

$$\left[\mathrm{Na}^+\right]_e = -\left[K^+\right]_e + \left[\mathrm{Cl}^-\right]_e - z_Y\left[Y^{z_Y}\right]_e \tag{A8b}$$

To achieve positive ion concentrations, i.e., $[Na^+]_e > 0$ and $[Cl^-]_e > 0$, we must have $[Cl^-]_e = 0.5\left(\mathcal{O}_e + (z_Y - 1)\left[Y^{z_Y}\right]_e\right) > [K^+]_e + z_Y[Y^{z_Y}]_e$, i.e., $\mathcal{O}_e > 2[K^+]_e + (1 + z_Y)[Y^{z_Y}]_e$.

### B. Stationary solutions of KJU systems

The PLEs for the KJU system, defined by $g_{K,1} = 0$ and $g_{Na,2} = 0$, and assuming $g_{Na,P} = 0$ (i.e., corresponding to the $[0, 1, 1]$ or $[0, 1, 0]$ configurations on the paracellular pathway), together with an NKA pump on the apical membrane, are given by the following equations for the ionic concentrations in compartments $A$ and $B$:

$$F\frac{d\,(w_A[Na^+]_A)}{dt} = -g_{Na,1}\,(V_A - E_{Na,A}) - \gamma_{Na} p a_2 \tag{B1a}$$

$$F\frac{d\,(w_A[K^+]_A)}{dt} = -g_{K,2}\,[(V_A - E_{K,A}) - (V_B - E_{K,B})] + \gamma_K p a_2 \tag{B1b}$$

$$F\frac{d\,(w_A[Cl^-]_A)}{dt} = g_{Cl,1}\,(V_A - E_{Cl,A}) + g_{Cl,2}\,[(V_A - E_{Cl,A}) - (V_B - E_{Cl,B})] \tag{B1c}$$

$$F\frac{d\,(w_B[Na^+]_B)}{dt} = \gamma_{Na} p a_2 \tag{B1d}$$

$$F\frac{d\,(w_B[K^+]_B)}{dt} = -g_{K,P}\,(V_B - E_{K,B}) + g_{K,2}\,[(V_A - E_{K,A}) - (V_B - E_{K,B})] - \gamma_K p a_2 \tag{B1e}$$

$$F\frac{d\,(w_B[Cl^-]_B)}{dt} = g_{Cl,P}\,(V_B - E_{Cl,B}) - g_{Cl,2}\,[(V_A - E_{Cl,A}) - (V_B - E_{Cl,B})] \tag{B1f}$$

At stationarity, the variables satisfy the following electroneutrality relations in compartments $A$ and $B$:

$$[Na^+]_A^{ss} + [K^+]_A^{ss} - [Cl^-]_A^{ss} + z_A\,[X^{z_A}]_A^{ss} = 0 \tag{B2a}$$

$$[[Na^+]_B^{ss} + [K^+]_B^{ss} - [Cl^-]_B^{ss} + z_B\,[X^{z_B}]_B^{ss} = 0 \tag{B2b}$$

In what follows, we seek a *stationary* solution of the PLEs in which all ionic concentrations in compartments $A$ and $B$, as well as the volume of compartment $A$, converge to constant steady states, while the volume of compartment $B$ grows linearly in time, say with slope $m_B > 0$.

To simplify the analysis, we further assume that $z_A = -1$. The results can be extended to more general parameter choices.

**Proposition 1** *Consider the PLEs described by Equations* (B1)*,* (A2)*,* (A3) *with the following conditions on the parameter values:*

*(i)* $z_A = -1$

*(ii) All conductances in the equations are positive, except for* $g_{Cl,P}$*, which is allowed to be nonnegative.*

*(iii)* $\frac{\mathcal{O}_e^2}{4(Na_e+K_e)} > Cl_e$.

*Then, (1) for any* $p > 0$*, there exists a corresponding* $m_B = m_B(p) > 0$ *for which* $w_B$ *grows linearly with rate* $m_B$*, while all other variables of the PLEs converge to steady-state values given by:*

$$[Na^+]_A^{ss} = \alpha Na_e e^{\frac{-F}{RT}\frac{\gamma_{Na}pa_2}{g_{Na,1}}} \tag{B3a}$$

$$[K^+]_A^{ss} = \alpha K_e e^{\frac{F}{RT}\left(\frac{\gamma_K pa_2}{g_{K,2}} - \frac{Fm_B}{g_{K,P}}\mathcal{A}\right)} \tag{B3b}$$

$$[Cl^-]_A^{ss} = \frac{Cl_e}{\alpha} e^{-\beta\left(\frac{\mathcal{O}_e}{2}+\frac{(\nu_1+\nu_2)m_B}{2S_\nu}\right)\frac{F^2 m_B}{RT}\frac{g_{Cl,2}}{g_{Cl,2}+g_{Cl,1}}} \tag{B3c}$$

$$[X]_A^{ss} = \frac{\mathcal{O}_e}{2} + \frac{\nu_2 m_B}{2S_\nu} - \frac{Cl_e}{\alpha} e^{-\beta\left(\frac{\mathcal{O}_e}{2}+\frac{(\nu_1+\nu_2)m_B}{2S_\nu}\right)\frac{F^2 m_B}{RT}\frac{g_{Cl,2}}{g_{Cl,2}+g_{Cl,1}}} \tag{B3d}$$

$$V_A^{ss} = -\frac{RT}{F}\ln\alpha \tag{B3e}$$

$$[Na^+]_B^{ss} = \frac{\gamma_{Na}pa_2}{Fm_B} \tag{B3f}$$

$$[K^+]_B^{ss} = \mathcal{A} \tag{B3g}$$

$$[Cl^-]_B^{ss} = \frac{\mathcal{O}_e}{2} + \frac{(\nu_1+\nu_2)m_B}{2S_\nu} \tag{B3h}$$

$$[X]_B^{ss} = 0 \tag{B3i}$$

$$V_B^{ss} = -\frac{RT}{F}\ln\frac{\mathcal{A}}{K_e} - \frac{Fm_B}{g_{K,P}}\mathcal{A} = E_{Cl,B}^{ss} + F\beta m_B [Cl]_B^{ss} \tag{B3j}$$

*where*

$$\mathcal{A} = \frac{\mathcal{O}_e}{2} + \frac{(\nu_1+\nu_2)m_B}{2S_\nu} - \frac{\gamma_{Na}pa_2}{Fm_B} \tag{B4a}$$

$$\alpha = \frac{\frac{\mathcal{O}_e}{2} + \frac{\nu_2 m_B}{2S_\nu}}{Na_e e^{\frac{-F}{RT}\frac{\gamma_{Na}pa_2}{g_{Na,1}}} + K_e e^{\frac{F}{RT}\left(\frac{\gamma_K pa_2}{g_{K,2}} - \frac{Fm_B}{g_{K,P}}\mathcal{A}\right)}} \tag{B4b}$$

$$\beta = \frac{g_{Cl,1}+g_{Cl,2}}{(g_{Cl,1}+g_{Cl,2})(g_{Cl,P}+g_{Cl,2}) - g_{Cl,2}^2} \tag{B4c}$$

$$S_\nu = \nu_1\nu_2 + \nu_1\nu_P + \nu_2\nu_P \tag{B4d}$$

*(2) There exists* $p_{\max} > 0$ *such that, whenever* $p \in (0, p_{\max})$*, all concentrations are non-negative.*

**Proof.** Since $[X^{z_B}]_B = \frac{X^{z_B}}{w_B}$, where the total amount $X^{z_B}$ of impermeant molecules in compartment $B$ is constant (i.e., $X^{z_B} = X^{z_B}(t=0)$), it follows that $[X^{z_B}]_B$ decreases as $w_B$ increases. In particular, if $w_B$ grows linearly in time, we have $[X^{z_B}]_B^{ss} = 0$.

To find the stationary solutions we let the RHS of Equations (B1a)–(B1c) and (A2a) equal to zero, the RHS of (A2b) equals to $m_B$, and using the chain rule, the RHS of Equations (B1d)–(B1f) equal to $Fm_B[\text{ion}]_B^{ss}$. Because

$$F\frac{d\,(w_B[\text{ion}]_B)}{dt} = F\frac{dw_B}{dt}[\text{ion}]_B + F\frac{d[\text{ion}]_B}{dt}w_B$$

which at the stationary solution, $\frac{d[\text{ion}]_B}{dt} = 0$, $\frac{dw_B}{dt} = m_B$ and $[\text{ion}]_B = [\text{ion}]_B^{ss}$.

Let $S_\nu = \nu_1\nu_2 + \nu_1\nu_P + \nu_2\nu_P$. Then, (A2a)= 0 and (A2b)= $m_B$ yield

$$[Na^+]_A^{ss} + [K^+]_A^{ss} + [Cl^-]_A^{ss} + [X^{z_A}]_A^{ss} = \mathcal{O}_e + \frac{\nu_2 m_B}{S_\nu} \tag{B5a}$$

$$[Na^+]_B^{ss} + [K^+]_B^{ss} + [Cl^-]_B^{ss} + [X^{z_B}]_B^{ss} = \mathcal{O}_e + \frac{(\nu_1+\nu_2)m_B}{S_\nu}. \tag{B5b}$$

The electroneutrality condition (B2a) and Equation (B5a) give the following relation for the stationary solution:

$$[Na^+]_A^{ss} + [K^+]_A^{ss} = \frac{1}{2}\left(\mathcal{O}_e + \frac{\nu_2 m_B}{S_\nu} - (1+z_A)\,[X^{z_A}]_A^{ss}\right) \tag{B6a}$$

$$[Cl^-]_A^{ss} = \frac{1}{2}\left(\mathcal{O}_e + \frac{\nu_2 m_B}{S_\nu} - (1-z_A)\,[X^{z_A}]_A^{ss}\right) \tag{B6b}$$

Similarly, the electroneutrality condition (B2b) and Equation (B5b) give:

$$[Na^+]_B^{ss} + [K^+]_B^{ss} = \frac{1}{2}\left(\mathcal{O}_e + \frac{(\nu_1+\nu_2)m_B}{S_\nu} - (1+z_B)\,[X^{z_B}]_B^{ss}\right)$$

$$[Cl^-]_B^{ss} = \frac{1}{2}\left(\mathcal{O}_e + \frac{(\nu_1+\nu_2)m_B}{S_\nu} - (1-z_B)\,[X^{z_B}]_B^{ss}\right)$$

The equality $[X^{z_B}]_B^{ss} = 0$ yields:

$$[Na]_B^{ss} + [K]_B^{ss} = \frac{\mathcal{O}_e}{2} + \frac{(\nu_1+\nu_2)m_B}{2S_\nu} \tag{B7a}$$

$$[Cl]_B^{ss} = \frac{\mathcal{O}_e}{2} + \frac{(\nu_1+\nu_2)m_B}{2S_\nu} \tag{B7b}$$

Equation (B1d)= $Fm_B[Na]_B^{ss}$ and Equation (B7) give

$$[Na^+]_B^{ss} = \frac{\gamma_{Na} p a_2}{Fm_B} \tag{B8a}$$

$$[K^+]_B = \frac{\mathcal{O}_e}{2} + \frac{(\nu_1+\nu_2)m_B}{2S_\nu} - \frac{\gamma_{Na} p a_2}{Fm_B} \tag{B8b}$$

$$[Cl^-]_B^{ss} = \frac{\mathcal{O}_e}{2} + \frac{(\nu_1+\nu_2)m_B}{2S_\nu} \tag{B8c}$$

Note that the sodium and chloride steady states are positive, assuming that all the parameter values are positive. However, to have a positive potassium concentration, we need the following condition on the parameter values

$$\frac{\mathcal{O}_e}{2}m_B + \frac{(\nu_1+\nu_2)}{2S_\nu}m_B^2 > \frac{\gamma_{Na}pa_2}{F}. \tag{B9}$$

Later, when we find the relationship between $m_B$ and $p$, we choose an $m_B$ that condition (B9) holds.

Next, we will derive an expression for $V_B^{ss}$. Note that $V_B^{ss}$ can be derived in two ways: (1) set (B1b)$=0$ and (B1e)$=Fm_B[K^+]_B^{ss}$ and use Equation (B8b):

$$V_B^{ss} = E_{K,B}^{ss} - \frac{Fm_B}{g_{K,P}}[K]_B^{ss} \tag{B10}$$

and (2) set (B1c)$=0$ to solve $V_A^{ss} - E_{Cl,A}^{ss}$ in terms of $V_B^{ss} - E_{Cl,B}^{ss}$ and substitute it into (B1f)$=Fm_B[Cl^-]_B^{ss}$:

$$V_B^{ss} = E_{Cl,B}^{ss} + F\beta m_B[Cl^-]_B^{ss} \tag{B11}$$

To have a well defined $V_B^{ss}$, we must equate Equations (B10)-(B11) which yield an implicit relation between $p$ and $m_B$. Later, we will show that this implicit relationship, gives an $m_B > 0$ for any fixed $p > 0$ such that (B9) holds. As long as we keep this relation between $p$ and $m_B$, we can choose any of these expressions for $V_B^{ss}$.

Next, we find the stationary values corresponding to compartment $A$. Equation (B1a)$=0$ gives the expression of $[Na^+]_A^{ss}$ in terms of $V_A^{ss}$

$$[Na^+]_A^{ss} = Na_e e^{\frac{-F}{RT}\left(\frac{\gamma_{Na}pa_2}{g_{Na,1}}+V_A^{ss}\right)}$$

Substituting $[K^+]_B^{ss}$ into Equation (B1b)$=0$ gives the expression of $[K^+]_A^{ss}$ in terms of $V_A^{ss}$

$$[K^+]_A^{ss} = K_e e^{\frac{F}{RT}\left(\frac{\gamma_K pa_2}{g_{K,2}}-\frac{Fm_B}{g_{K,P}}\left(\frac{\mathcal{O}_e}{2}+\frac{(\nu_1+\nu_2)m_B}{2S_\nu}-\frac{\gamma_{Na}pa_2}{Fm_B}\right)-V_A^{ss}\right)}$$

Now by adding these expressions for sodium and potassium and using Equation (C5a) with $z_A = -1$, we can get an expression for $V_A^{ss}$ as follows:

$$\left(Na_e e^{\frac{-F}{RT}\frac{\gamma_{Na}pa_2}{g_{Na,1}}} + K_e e^{\frac{F}{RT}\left(\frac{\gamma_K pa_2}{g_{K,2}}-\frac{Fm_B}{g_{K,P}}\left(\frac{\mathcal{O}_e}{2}+\frac{(\nu_1+\nu_2)m_B}{2S_\nu}-\frac{\gamma_{Na}pa_2}{Fm_B}\right)\right)}\right)e^{\frac{-F}{RT}V_A^{ss}}$$
$$= \frac{\mathcal{O}_e}{2} + \frac{\nu_2 m_B}{2S_\nu}$$

which yields to an explicit formula for $e^{\frac{-F}{RT}V_A^{ss}}$, that is, $e^{\frac{-F}{RT}V_A^{ss}} = \alpha$, and subsequently for $[Na^+]_A^{ss}$, $[K^+]_A^{ss}$, and $V_A^{ss}$.

Finally, similar to those in compartment $B$, we can derive $[Cl^-]_A^{ss}$ and use Equation (C5b) with $z_A = -1$ to derive $[X]_A^{ss}$ or equivalently $w_A^{ss}$.

To prove that for any $p > 0$ there exists an $m_B > 0$, we fix $p > 0$, equate (B10) and (B11):

$$\begin{aligned}
&\frac{RT}{F}\ln\left(\frac{\mathcal{O}_e}{2Cl_e} + \frac{(\nu_1+\nu_2)m_B}{2S_\nu Cl_e}\right) + Fm_B\beta\left(\frac{\mathcal{O}_e}{2} + \frac{(\nu_1+\nu_2)m_B}{2S_\nu}\right) \\
&= -\frac{RT}{F}\ln\left(\frac{\mathcal{O}_e}{2K_e} + \frac{(\nu_1+\nu_2)m_B}{2S_\nu K_e} - \frac{\gamma_{Na}pa_2}{Fm_BK_e}\right) \\
&\quad -\frac{Fm_B}{g_{K,P}}\left(\frac{\mathcal{O}_e}{2} + \frac{(\nu_1+\nu_2)m_B}{2S_\nu} - \frac{\gamma_{Na}pa_2}{Fm_B}\right)
\end{aligned}$$

which is equivalent to

$$\begin{aligned}
&\frac{RT}{F}\ln\left(\frac{\mathcal{O}_e}{2K_e} + \frac{(\nu_1+\nu_2)m_B}{2S_\nu K_e} - \frac{\gamma_{Na}pa_2}{Fm_BK_e}\right) + \frac{RT}{F}\ln\left(\frac{\mathcal{O}_e}{2Cl_e} + \frac{(\nu_1+\nu_2)m_B}{2S_\nu Cl_e}\right) \\
&= -\frac{Fm_B}{g_{K,P}}\left(\frac{\mathcal{O}_e}{2} + \frac{(\nu_1+\nu_2)m_B}{2S_\nu} - \frac{\gamma_{Na}pa_2}{Fm_B}\right) - Fm_B\beta\left(\frac{\mathcal{O}_e}{2} + \frac{(\nu_1+\nu_2)m_B}{2S_\nu}\right)
\end{aligned}$$

Multiplying both sides by $\frac{F}{RT}$, taking the exponential, and multiplying by $K_eCl_em_B$ we gain

$$\mathcal{F}(m_B) = \mathcal{G}(m_B)$$

where

$$\mathcal{F}(m_B) = \left(\frac{\mathcal{O}_e}{2}m_B + \frac{(\nu_1+\nu_2)m_B^2}{2S_\nu} - \frac{\gamma_{Na}pa_2}{F}\right)\left(\frac{\mathcal{O}_e}{2} + \frac{(\nu_1+\nu_2)m_B}{2S_\nu}\right)$$

and

$$\mathcal{G}(m_B) = K_eCl_em_Be^{\frac{F\gamma_{Na}a_2}{RTg_{K,P}}p}e^{-\frac{F^2}{RT}(\frac{1}{g_{K,P}}+\beta)m_B\left(\frac{\mathcal{O}_e}{2}+\frac{(\nu_1+\nu_2)m_B}{2S_\nu}\right)}$$

We show that $\mathcal{F}$ and $\mathcal{G}$ intersect at some $m_B$ where

$$m_B > m_B^* := \frac{\frac{-\mathcal{O}_e}{2} + \sqrt{\frac{\mathcal{O}_e^2}{4} + \frac{2(\nu_1+\nu_2)\gamma_{Na}pa_2}{FS_\nu}}}{\frac{\nu_1+\nu_2}{S_\nu}} > 0$$

Note that $\mathcal{F}(m_B^*) = 0$ and $\mathcal{F}(m_B) \to \infty$ as $m_B \to \infty$. On the other hand, $\mathcal{G}(m_B^*) > 0$ and $\mathcal{G}(m_B) \to 0$ as $m_B \to \infty$. Therefore, since $\mathcal{F}$ and $\mathcal{G}$ are continuous, there exists an $m_B > m_B^*$ such that $\mathcal{F}(m_B) = \mathcal{G}(m_B)$.

Note that all concentrations are naturally non-negative except for $[X]_A^{ss}$. In what follows, we use condition (iii) of the proposition to find an interval for $p$ in which $[X]_A^{ss} > 0$. Let

$$f(p, m_B) := \alpha\ [X]_A^{ss} = \alpha(p, m_B)\left(\frac{\mathcal{O}_e}{2} + \frac{\nu_2 m_B(p)}{2S_\nu}\right) - Cl_e\ e^{-\beta\left(\frac{\mathcal{O}_e}{2} + \frac{(\nu_1+\nu_2)m_B}{2S_\nu}\right)\frac{F^2 m_B}{RT}\frac{g_{Cl,2}}{g_{Cl,2}+g_{Cl,1}}}$$

Then, the positivity of $[X]_A^{ss}$ is equivalent to $f(p, m_B) > 0$. Since $f$ is increasing in $m_B$ and $m_B > m_B^*$, it is sufficient to show that $f(p, m_B^*) > 0$ for some $p > 0$. Define

$$g(p) := f(p, m_B^*(p)) > 0.$$

Then $g(p) > 0$ implies $f(p, m_B) > 0$ for all $m_B > m_B^*$. Since $m_B^*(0) = 0$, we have

$$g(0) = \frac{\mathcal{O}_e^2}{4(Na_e + K_e)} - Cl_e > 0 \quad \text{(by assumption (iii) of the proposition).}$$

Therefore, either $g(p) > 0$ for any $p > 0$, i.e., $p_{\max} = \infty$ or there exists $0 < p_{\max} < \infty$ such that $g(p_{\max}) = 0$ and $g(p) > 0$ for $p \in (0, p_{\max})$. Hence, $[X]_A^{ss} > 0$ for $p \in (0, p_{\max})$. Note that $g$ might have multiple zeros but we are interested in its smallest zero and call it $p_{\max}$.

**Remark (Necessary conditions on ionic conductances for stability in $A$ and linear expansion in $B$)** Consider an active KJU system with $p > 0$ on side 2, and $g_{\text{Na},2} = g_{\text{K},1} = 0$. Suppose that we seek a stationary state characterized by positive steady-state ion concentrations, a positive steady-state volume $A$, and a linear-type stationary state for volume $B$ with positive slope $m_B > 0$. In other words, we impose $p > 0, w_A^{\text{ss}} > 0, m_B > 0, [\text{ion}]_{A,B}^{\text{ss}} > 0$. For such a stationary state to exist, the conductances must satisfy the following conditions:

$$g_{\text{Na},1} > 0, \quad g_{\text{K},2} > 0, \quad g_{\text{K},P} > 0, \quad g_{\text{Cl},P} > 0,$$

or, alternatively,

$$g_{\text{Cl},P} = 0, \quad \text{and} \quad g_{\text{Cl},1} \times g_{\text{Cl},2} > 0.$$

Equivalently, the chloride condition can be written compactly as

$$g_{\text{Cl},P} > 0 \quad \text{or} \quad g_{\text{Cl},P} = 0 \text{ and } g_{\text{Cl},1} \times g_{\text{Cl},2} > 0.$$

### C. Stationary solutions of reverse KJU systems

The PLEs for the reverse KJU (rKJU) system, defined by $g_{K,1} = 0$ and $g_{Na,2} = 0$, and assuming $g_{K,P} = 0$ (i.e., corresponding to the $[1, 0, 1]$ or $[1, 0, 0]$ configurations on the paracellular pathway), together with an NKA pump on the basolateral membrane, are given by the following equations for the ionic concentrations in compartments $A$ and $B$:

$$F\frac{d\left(w_A[Na]_A\right)}{dt} = -g_{Na,1}\left(V_A - E_{Na,A}\right) - \gamma_{Na} p a_1 \tag{C1a}$$

$$F\frac{d\left(w_A[K]_A\right)}{dt} = -g_{K,2}\left[\left(V_A - E_{K,A}\right) - \left(V_B - E_{K,B}\right)\right] + \gamma_K p a_1 \tag{C1b}$$

$$F\frac{d\left(w_A[Cl]_A\right)}{dt} = g_{Cl,1}\left(V_A - E_{Cl,A}\right) + g_{Cl,2}\left[\left(V_A - E_{Cl,A}\right) - \left(V_B - E_{Cl,B}\right)\right] \tag{C1c}$$

$$F\frac{d\left(w_B[Na]_B\right)}{dt} = -g_{Na,P}\left(V_B - E_{Na,B}\right) \tag{C1d}$$

$$F\frac{d\left(w_B[K]_B\right)}{dt} = g_{K,2}\left[\left(V_A - E_{K,A}\right) - \left(V_B - E_{K,B}\right)\right] \tag{C1e}$$

$$F\frac{d\left(w_B[Cl]_B\right)}{dt} = g_{Cl,P}\left(V_B - E_{Cl,B}\right) - g_{Cl,2}\left[\left(V_A - E_{Cl,A}\right) - \left(V_B - E_{Cl,B}\right)\right] \tag{C1f}$$

Similar to the KJU systems, the volumes and membrane potentials of compartments $A$ and $B$ evolve according to Equations (A2) and by(A3). Also, the variables satisfy the electroneutrality relations in (B2).

In what follows, we seek a *stationary* solution of the rKJU system in which all ionic concentrations in compartments $A$ and $B$, as well as the volume of compartment $A$, converge to constant steady states, while the volume of compartment $B$ grows linearly in time, say with slope $m_B > 0$.

To simplify the analysis, we further assume that $z_A = -1$. The results can be extended to more general parameter choices.

**Proposition 2** *Consider the rKJU described by Equations* (A2), (A3), *and* (C1) *with the following conditions on the parameter values:*

*(i)* $z_A = -1$

*(ii) All conductances in the equations are positive, except for* $g_{Cl,P}$, *which is allowed to be nonnegative.*

*(iii)* $\frac{\mathcal{O}_e^2}{4Na_e} > Cl_e$.

*Then, (1) for any* $p > 0$, *there exists a corresponding* $m_B = m_B(p) > 0$ *for which* $w_B$ *grows linearly with rate* $m_B$, *while all other variables of the*

*PLEs converge to steady-state values given by:*

$$[Na^+]_A^{ss} = \alpha Na_e e^{\frac{-F}{RT}\frac{\gamma_{Na} pa_1}{g_{Na,1}}} \tag{C2a}$$

$$[K^+]_A^{ss} = \frac{\mathcal{O}_e}{2} + \frac{\nu_2 m_B}{2S_\nu} - [Na]_A^{ss} \tag{C2b}$$

$$[Cl^-]_A^{ss} = \frac{Cl_e}{\alpha} e^{-\beta\left(\frac{\mathcal{O}_e}{2}+\frac{(\nu_1+\nu_2)m_B}{2S_\nu}\right)\frac{F^2 m_B}{RT}\frac{g_{Cl,2}}{g_{Cl,2}+g_{Cl,1}}} \tag{C2c}$$

$$[X]_A^{ss} = \frac{\mathcal{O}_e}{2} + \frac{\nu_2 m_B}{2S_\nu} - \frac{Cl_e}{\alpha} e^{-\beta\left(\frac{\mathcal{O}_e}{2}+\frac{(\nu_1+\nu_2)m_B}{2S_\nu}\right)\frac{F^2 m_B}{RT}\frac{g_{Cl,2}}{g_{Cl,2}+g_{Cl,1}}} \tag{C2d}$$

$$V_A^{ss} = -\frac{RT}{F}\ln\alpha \tag{C2e}$$

$$[Na^+]_B^{ss} = \mathcal{A} \tag{C2f}$$

$$[K^+]_B^{ss} = \frac{\gamma_K pa_1}{F m_B} \tag{C2g}$$

$$[Cl^-]_B^{ss} = \frac{\mathcal{O}_e}{2} + \frac{(\nu_1+\nu_2)m_B}{2S_\nu} \tag{C2h}$$

$$[X]_B^{ss} = 0 \tag{C2i}$$

$$V_B^{ss} = -\frac{F m_B}{g_{Na,P}}\mathcal{A} - \frac{RT}{F}\ln\frac{\mathcal{A}}{Na_e} \tag{C2j}$$

*where*

$$\mathcal{A} = \frac{\mathcal{O}_e}{2} + \frac{(\nu_1+\nu_2)m_B}{2S_\nu} - \frac{\gamma_K pa_1}{F m_B} \tag{C3a}$$

$$\alpha = \frac{\frac{\mathcal{O}_e}{2} + \frac{\nu_2 m_B}{2S_\nu}}{Na_e e^{\frac{-F}{RT}\frac{\gamma_{Na} pa_1}{g_{Na,1}}} + \frac{\gamma_K pa_1}{F m_B}\frac{Na_e}{\mathcal{A}} e^{\frac{F}{RT}\left(\frac{\gamma_K pa_1}{g_{K,2}} - \frac{F m_B}{g_{Na,P}}\mathcal{A}\right)}} \tag{C3b}$$

$$\beta = \frac{g_{Cl,1}+g_{Cl,2}}{(g_{Cl,1}+g_{Cl,2})(g_{Cl,P}+g_{Cl,2}) - g_{Cl,2}^2} \tag{C3c}$$

$$S_\nu = \nu_1\nu_2 + \nu_1\nu_P + \nu_2\nu_P \tag{C3d}$$

*(2) There exists $p_{\max} > 0$ such that, whenever $p \in (0, p_{\max})$, all concentrations are non-negative.*

**Proof.** Since $[X^{z_B}]_B = \frac{X^{z_B}}{w_B}$, where the total amount $X^{z_B}$ of impermeant molecules in compartment $B$ is constant (i.e., $X^{z_B} = X^{z_B}(t=0)$), it follows that $[X^{z_B}]_B$ decreases as $w_B$ increases. In particular, if $w_B$ grows linearly in time, we have $[X^{z_B}]_B^{ss} = 0$.

To find the stationary solutions we let the RHS of Equations (C1a)–(C1c) and (A2a) equal to zero, the RHS of (A2b) equals to $m_B$, and using the chain rule, the RHS of Equations (C1d)–(C1f) equal to $Fm_B[\text{ion}]_B^{ss}$. Because

$$F\frac{d\left(w_B[\text{ion}]_B\right)}{dt} = F\frac{dw_B}{dt}[\text{ion}]_B + F\frac{d[\text{ion}]_B}{dt}w_B$$

which at the stationary solution, $\frac{d[\text{ion}]_B}{dt} = 0$, $\frac{dw_B}{dt} = m_B$ and $[\text{ion}]_B = [\text{ion}]_B^{ss}$.

Let $S_\nu = \nu_1\nu_2 + \nu_1\nu_P + \nu_2\nu_P$. Then, (A2a)$= 0$ and (A2b)$= m_B$ yield

$$[Na^+]_A^{ss} + [K^+]_A^{ss} + [Cl^-]_A^{ss} + [X^{z_A}]_A^{ss} = \mathcal{O}_e + \frac{\nu_2 m_B}{S_\nu} \tag{C4a}$$

$$[Na^+]_B^{ss} + [K^+]_B^{ss} + [Cl^-]_B^{ss} + [X^{z_B}]_B^{ss} = \mathcal{O}_e + \frac{(\nu_1+\nu_2) m_B}{S_\nu}. \tag{C4b}$$

The electroneutrality condition (B2a) and Equation (C4a) give the following relation for the stationary solution:

$$[Na^+]_A^{ss} + [K^+]_A^{ss} = \frac{1}{2}\left(\mathcal{O}_e + \frac{\nu_2 m_B}{S_\nu} - (1+z_A)\,[X^{z_A}]_A^{ss}\right) \tag{C5a}$$

$$[Cl^-]_A^{ss} = \frac{1}{2}\left(\mathcal{O}_e + \frac{\nu_2 m_B}{S_\nu} - (1-z_A)\,[X^{z_A}]_A^{ss}\right) \tag{C5b}$$

Similarly, the electroneutrality condition (B2b) and Equation (C4b) give:

$$[Na^+]_B^{ss} + [K^+]_B^{ss} = \frac{1}{2}\left(\mathcal{O}_e + \frac{(\nu_1+\nu_2) m_B}{S_\nu} - (1+z_B)\,[X^{z_B}]_B^{ss}\right)$$

$$[Cl^-]_B^{ss} = \frac{1}{2}\left(\mathcal{O}_e + \frac{(\nu_1+\nu_2) m_B}{S_\nu} - (1-z_B)\,[X^{z_B}]_B^{ss}\right)$$

The equality $[X^{z_B}]_B^{ss} = 0$ yields:

$$[Na^+]_B^{ss} + [K^+]_B^{ss} = \frac{\mathcal{O}_e}{2} + \frac{(\nu_1+\nu_2) m_B}{2S_\nu} \tag{C6a}$$

$$[Cl^-]_B^{ss} = \frac{\mathcal{O}_e}{2} + \frac{(\nu_1+\nu_2) m_B}{2S_\nu} \tag{C6b}$$

Plugging Equation (C1b)$= 0$ into Equation (C1e)$= Fm_B[K]_B^{ss}$ and Equation (C6) give

$$[K^+]_B^{ss} = \frac{\gamma_K p a_1}{Fm_B} \tag{C7a}$$

$$[Na^+]_B^{ss} = \frac{\mathcal{O}_e}{2} + \frac{(\nu_1+\nu_2) m_B}{2S_\nu} - \frac{\gamma_K p a_1}{Fm_B} \tag{C7b}$$

$$[Cl^-]_B^{ss} = \frac{\mathcal{O}_e}{2} + \frac{(\nu_1+\nu_2) m_B}{2S_\nu} \tag{C7c}$$

Note that the potassium and chloride steady states are positive, assuming that all the parameter values are positive. However, to have a positive sodium concentration, we need the following condition on the parameter values

$$\frac{\mathcal{O}_e}{2} m_B + \frac{(\nu_1 + \nu_2)}{2S_\nu} m_B^2 > \frac{\gamma_K p a_1}{F}. \tag{C8}$$

Later, when we find the relationship between $m_B$ and $p$, we choose an $m_B$ that condition (C8) holds.

Next, we will derive an expression for $V_B^{ss}$. Note that $V_B^{ss}$ can be derived in two ways: (i) set (C1d)$= F m_B [Na^+]_B^{ss}$ and use Equation (C7a):

$$V_B^{ss} = E_{Na,B}^{ss} - \frac{F m_B}{g_{Na,P}} [Na^+]_B^{ss} \tag{C9}$$

and (ii) set (C1c)$= 0$ to solve $V_A^{ss} - E_{Cl,A}^{ss}$ in terms of $V_B^{ss} - E_{Cl,B}^{ss}$ and substitute it into (C1f)$= F m_B [Cl^-]_B^{ss}$:

$$V_B^{ss} = E_{Cl,B}^{ss} + F \beta m_B [Cl^-]_B^{ss} \tag{C10}$$

To have a well defined $V_B^{ss}$, we must equate Equations (C9)-(C10) which yield an implicit relation between $p$ and $m_B$. Later, we will show that this implicit relationship, gives an $m_B > 0$ for any fixed $p > 0$ such that (B9) holds. As long as we keep this relation between $p$ and $m_B$, we can choose any of these expressions for $V_B^{ss}$.

Next, we find the stationary values corresponding to compartment $A$. Equation (C1a)$= 0$ gives the expression of $[Na^+]_A^{ss}$ in terms of $V_A^{ss}$

$$[Na^+]_A^{ss} = Na_e e^{\frac{-F}{RT}\left(\frac{\gamma_{Na} p a_1}{g_{Na,1}} + V_A^{ss}\right)}$$

Substituting $V_B^{ss}$ and $[K^+]_B^{ss}$ into Equation (C1b)$= 0$ gives the expression of $[K^+]_A^{ss}$ in terms of $V_A^{ss}$

$$[K^+]_A^{ss} = K_e e^{\frac{-F}{RT} V_A^{ss}} e^{\frac{F}{RT}\left(\frac{\gamma_K p a_1}{g_{K,2}} + V_B^{ss} - E_{K,B}^{ss}\right)}$$

Now by adding these expressions for sodium and potassium and using Equation (C5a) with $z_A = -1$, we can get an expression for $V_A^{ss}$ as follows:

$$\left( Na_e e^{\frac{-F}{RT}\frac{\gamma_{Na} p a_1}{g_{Na,1}}} + K_e e^{\frac{F}{RT}\left(\frac{\gamma_K p a_1}{g_{K,2}} + V_B^{ss} - E_{K,B}^{ss}\right)} \right) e^{\frac{-F}{RT} V_A^{ss}} = \frac{\mathcal{O}_e}{2} + \frac{\nu_2 m_B}{2S_\nu}$$

which yields to an explicit formula for $e^{\frac{-F}{RT}V_A^{ss}}$, that is, $e^{\frac{-F}{RT}V_A^{ss}} = \alpha$, and subsequently for $[Na^+]_A^{ss}$, $[K^+]_A^{ss}$, and $V_A^{ss}$.

Finally, similar to those in compartment $B$, we can derive $[Cl]_A^{ss}$ and use Equation (C5b) with $z_A = -1$ to derive $[X]_A^{ss}$ or equivalently $w_A^{ss}$.

To prove that for any $p > 0$ there exists an $m_B > 0$, we fix $p > 0$, equate (C9) and (C10):

$$\begin{aligned}&\frac{RT}{F}\ln\left(\frac{\mathcal{O}_e}{2Cl_e}+\frac{(\nu_1+\nu_2)m_B}{2S_\nu Cl_e}\right)+Fm_B\beta\left(\frac{\mathcal{O}_e}{2}+\frac{(\nu_1+\nu_2)m_B}{2S_\nu}\right)\\&=-\frac{RT}{F}\ln\left(\frac{\mathcal{O}_e}{2Na_e}+\frac{(\nu_1+\nu_2)m_B}{2S_\nu Na_e}-\frac{\gamma_K pa_1}{Fm_BNa_e}\right)\\&\quad-\frac{Fm_B}{g_{Na,P}}\left(\frac{\mathcal{O}_e}{2}+\frac{(\nu_1+\nu_2)m_B}{2S_\nu}-\frac{\gamma_K pa_1}{Fm_B}\right)\end{aligned}$$

which is equivalent to

$$\begin{aligned}&\frac{RT}{F}\ln\left(\frac{\mathcal{O}_e}{2Na_e}+\frac{(\nu_1+\nu_2)m_B}{2S_\nu Na_e}-\frac{\gamma_K pa_1}{Fm_BNa_e}\right)+\frac{RT}{F}\ln\left(\frac{\mathcal{O}_e}{2Cl_e}+\frac{(\nu_1+\nu_2)m_B}{2S_\nu Cl_e}\right)\\&=-\frac{Fm_B}{g_{Na,P}}\left(\frac{\mathcal{O}_e}{2}+\frac{(\nu_1+\nu_2)m_B}{2S_\nu}-\frac{\gamma_K pa_1}{Fm_B}\right)-Fm_B\beta\left(\frac{\mathcal{O}_e}{2}+\frac{(\nu_1+\nu_2)m_B}{2S_\nu}\right)\end{aligned}$$

Multiplying both sides by $\frac{F}{RT}$, taking the exponential, and multiplying by $Na_eCl_em_B$ we gain

$$\mathcal{F}(m_B) = \mathcal{G}(m_B)$$

where

$$\mathcal{F}(m_B) = \left(\frac{\mathcal{O}_e}{2}m_B + \frac{(\nu_1+\nu_2)m_B^2}{2S_\nu} - \frac{\gamma_K pa_1}{F}\right)\left(\frac{\mathcal{O}_e}{2}+\frac{(\nu_1+\nu_2)m_B}{2S_\nu}\right)$$

and

$$\mathcal{G}(m_B) = Na_eCl_em_Be^{\frac{F\gamma_K a_1}{RTg_{Na,P}}p}e^{-\frac{F^2}{RT}(\frac{1}{g_{Na,P}}+\beta)m_B\left(\frac{\mathcal{O}_e}{2}+\frac{(\nu_1+\nu_2)m_B}{2S_\nu}\right)}$$

We show that $\mathcal{F}$ and $\mathcal{G}$ intersect at some $m_B$ where

$$m_B > m_B^* := \frac{\frac{-\mathcal{O}_e}{2}+\sqrt{\frac{\mathcal{O}_e^2}{4}+\frac{2(\nu_1+\nu_2)\gamma_K pa_1}{FS_\nu}}}{\frac{\nu_1+\nu_2}{S_\nu}} > 0$$

Note that $\mathcal{F}(m_B^*) = 0$ and $\mathcal{F}(m_B) \to \infty$ as $m_B \to \infty$. On the other hand, $\mathcal{G}(m_B^*) > 0$ and $\mathcal{G}(m_B) \to 0$ as $m_B \to \infty$. Therefore, since $\mathcal{F}$ and $\mathcal{G}$ are continuous, there exists an $m_B > m_B^*$ such that $\mathcal{F}(m_B) = \mathcal{G}(m_B)$.

The proof of part (2) is identical to that of Proposition 1.
